# A *Plasmodium falciparum* Pangenome Resource to drive Structural Variant Discovery and assist Malaria Control

**DOI:** 10.64898/2026.05.19.726271

**Authors:** Nina Billows, Joseph Thorpe, Susana Campino, Taane G. Clark

**Author notes:** Corresponding Author: Professor Taane Clark. Joint authors, Prof. Susana Campino, LSHTM Prof. Taane G. Clark, LSHTM.

## Abstract

*Plasmodium falciparum*, the deadliest causative agent of malaria, harbours extensive structural variation that underlies key biological processes including drug resistance and diagnostic evasion. Here, we present PfPan, a *P. falciparum* pangenome constructed from 13 geographically diverse high-quality reference genomes using Minigraph-Cactus, adding 4.77 Mb of sequence beyond the 3D7 linear reference. We identify over 5,000 structural variants (SVs) across the reference genomes and demonstrate improved genotyping using the vg toolkit compared to linear reference-based approaches, with comparable performance for small variant discovery. Benchmarking against assembly-derived truth sets confirms pangenome superiority for structural variant detection, particularly at complex and hypervariable loci. Applying PfPan to 878 globally sampled *P. falciparum* whole-genome sequences, we characterise biologically and clinically relevant SVs, including the complex drug resistance-linked *gch1* locus, and explore deletions upstream of the coding sequence for the diagnostic target HRP2. PfPan provides a foundational resource for mitigating reference bias in *P. falciparum* genomic surveillance and establishes a framework for improved detection of variants associated with drug resistance and malaria control in monoclonal *P. falciparum* samples that have not undergone selective whole-genome amplification.

## INTRODUCTION

Over two decades ago, the first reference genome for *Plasmodium falciparum,* the ‘deadliest’ causative agent of malaria, was published ^1^. The reference genome was constructed using isolate *P. falciparum* 3D7, a clone derived from NF54, a strain with predicted African origin that was originally isolated from a patient who had ‘airport malaria’ in the Netherlands ^2^. Since its original publication in 2002, the quality of the 3D7 reference genome assembly and its associated annotation have increased, providing a valuable resource for genomic analysis of *P. falciparum* ^3^. Whilst the 3D7 reference genome has continued to undergo significant improvement and many genomic studies have utilised it, there are several limitations associated with its use.

It is widely acknowledged that *P. falciparum* populations exhibit extensive genetic diversity and geographic patterns of genetic variation ^4^. Therefore, although the 3D7 reference genome is accurate, it is not entirely representative of global parasite populations, and genomic regions that are drastically divergent may not map well to a single reference genome, leading to ‘reference bias’. This concept is well recognised, and often genes that are highly variable and polymorphic, such as *var, rifin* and *stevor,* are excluded from population genomic analyses due to known mapping errors ^5^. In addition, variation in genes that are absent in 3D7 will also not be captured using a single reference ^6^. These limitations highlight that a single reference genome cannot fully capture the breadth of genetic variation present across *P. falciparum* populations.

After recognising that the use of a single reference genome cannot wholly represent global *P. falciparum* variation, a set of reference genomes from different populations was curated. This panel consists of near-complete, long-read assemblies for 15 *P. falciparum* isolates, including ten cultured clinical isolates ^6^. This dataset has enabled a clearer definition of the conserved core genome and subtelomeric boundaries, as well as analyses of structural variation and gene family diversity ^7^. However, its use has been constrained by the predominance of bioinformatic pipelines designed around a single reference genome.

To overcome reference bias and use diverse reference genome assemblies, “pangenome” references have been proposed to improve mapping and variant calling. For example, compared to the existing reference GRCh38, a draft human pangenome released in 2023 incorporates 119 million base pairs of euchromatic polymorphic sequences and 1,115 gene duplications ^8^. Whilst enhancing the detection of structural variants (SVs), the human pangenome also reduced small variant calling errors. Pangenomes have also been constructed for a diverse range of eukaryotic species, including yeast, plants, chickens and livestock, with similar advancements reported ^9–13^. Previously, ‘genome graphs’ have been applied to *P. falciparum* and successfully revealed previously hidden recombination ^14^. However, a complete pangenome reference is yet to be released and will require the adoption of new bioinformatic pipelines to aid processing. The identification of SVs is important for *P. falciparum,* with copy number variants, such as those in *mdr1*, being implicated in drug resistance ^4,15^. In addition, deletions in the histidine-rich protein-encoding genes (*hrp2/3*) are an important adaptive strategy to evade diagnostic detection. To date, detection of such variation from short-read data using established methods has remained challenging ^15^.

Here, we develop the first draft pangenome reference for *P. falciparum* (PfPan) and showcase the advantages of its use for small variant calling and structural variant discovery. We construct PfPan using the existing 3D7 reference (version 3) and long-read assemblies ^6^. We subsequently use this resource as a backbone to map 878 geographically diverse monoclonal *P. falciparum* isolates from publicly available short-read sequences. By comparing our approach with traditional linear mapping methods based on the 3D7 reference genome, we demonstrate the advantages of a pangenome-based approach.

## RESULTS

### Composition of the *Plasmodium falciparum* Pangenome

The *P. falciparum* pangenome (PfPan) was constructed using 3D7 and 12 high-quality, near-complete reference genomes from different geographic regions (**Table 1**). The initial selection comprised 3D7 and 15 additional strains with assembled genomes; three were subsequently excluded owing to polyclonal origin (TG01 and ML01) and assembly inconsistencies (SD01)^6^. All but one strain (CD01) had apicoplast and mitochondrial assemblies, in addition to the 14 nuclear chromosomes. Although the assemblies selected were near-complete, only two strains, 3D7 and GN01, had telomeres for all 14 chromosomes. To improve read mappability, we generated clipped and filtered graphs by removing large unaligned sequence segments and nodes unique to a single assembly. The final graph spanned 28,064,061 bp and comprised 1,398,468 nodes connected by 2,019,188 edges. The inclusion of 12 strain reference genomes alongside 3D7 added approximately 4.77 Mb of sequence compared with using 3D7 alone (**Figure 1A-C**). Despite this increase, most of the *P. falciparum* genome is conserved across all 13 haplotypes, forming a core backbone. Consistent with this, the pangenome graph exhibited a mean node degree of 2.89, close to the baseline of 2 expected for a fully linear graph. This indicates that the parasite genome is largely collinear across strains at the sequence level, with most nodes representing shared, non-branching sequence, while structural complexity is confined to a relatively small number of highly variable regions, such as subtelomeric and multigene family loci (**Supplementary Figure 1**). As the most complete assembly, 3D7 contributed the largest proportion of unique sequence to the graph (5.82%), whereas each of the remaining samples contributed less than 2%. (**Table 1**). Visualisation of the pangenome confirmed the existing, established boundaries of subtelomeric and internal hypervariable regions (**Supplementary Figure 2**).

**Figure 1.**
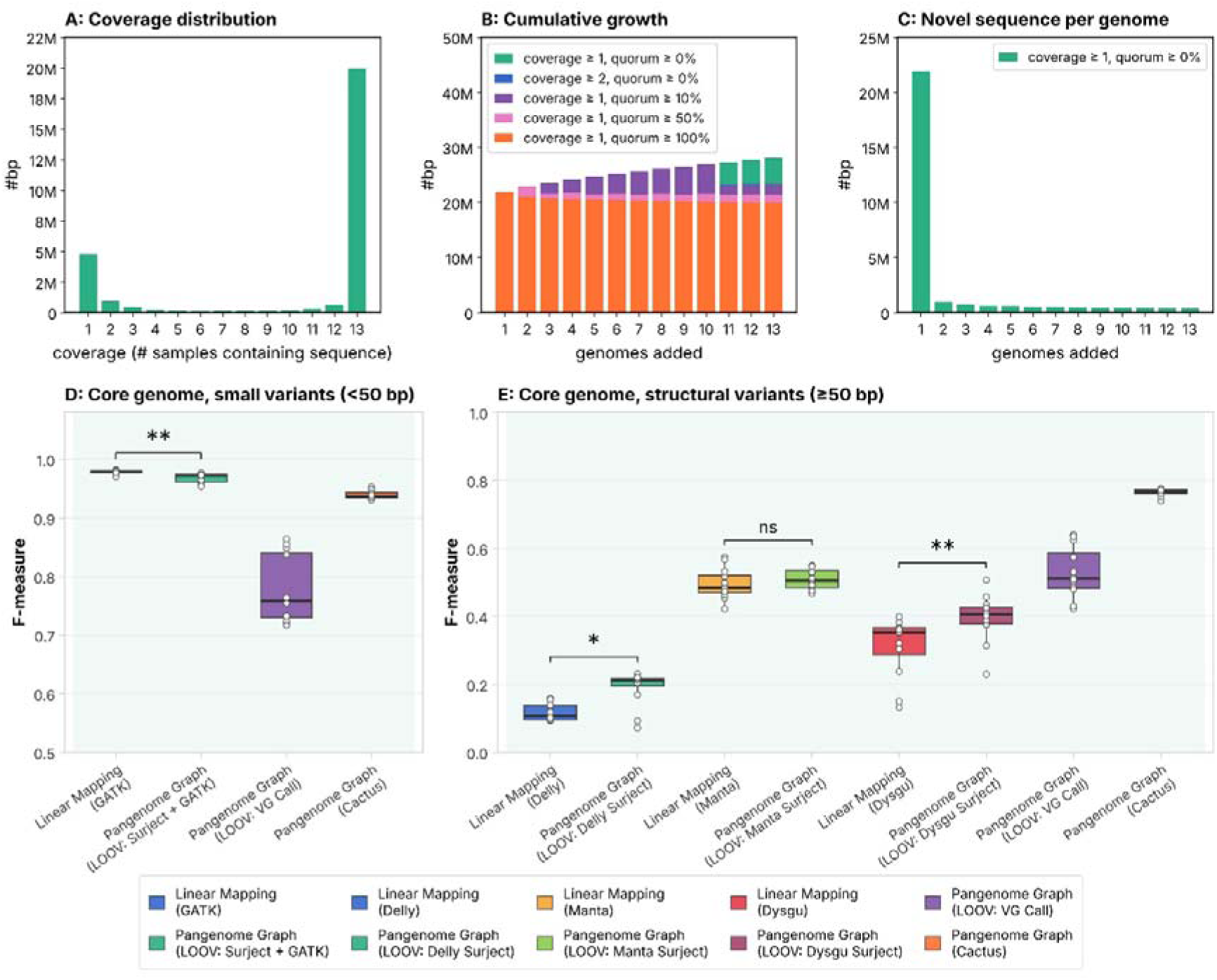
PfPan Growth Statistics and Variant Call Benchmarking. **(A)** Distribution of base-pair coverage: the amount of sequence (bp) present in exactly *n* of the 13 genomes (n = 1, private; n = 13, core). **(B)** Cumulative pangenome size (bp) as genomes are added one at a time, shown across five coverage/quorum thresholds. **(C)** Novel sequence (bp) contributed by each additional genome. **(D)** F-measure for small variants (<50 bp) and **(E)** structural variants (SVs) (≥50 bp) in the core genome, across the 12 strains tested by leave-one-out validation (LOOV; the graph is rebuilt without the tested strain). Linear Mapping and Pangenome Graph (LOOV: Surject) approaches call variants *de novo* from sequencing reads, aligned either to the linear reference or to the graph and then surjected onto it. Pangenome Graph (LOOV: VG Call) uses *genotyping only*: assigns genotypes to variants already present in the graph and cannot discover new ones. Brackets show two-sided paired Wilcoxon signed-rank tests of each linear caller against its graph-surject counterpart (Benjamini–Hochberg adjusted; *p < 0.05, **p < 0.01, ns not significant). Boxes show the median and interquartile range; each dot is one strain.

**Table 1.**
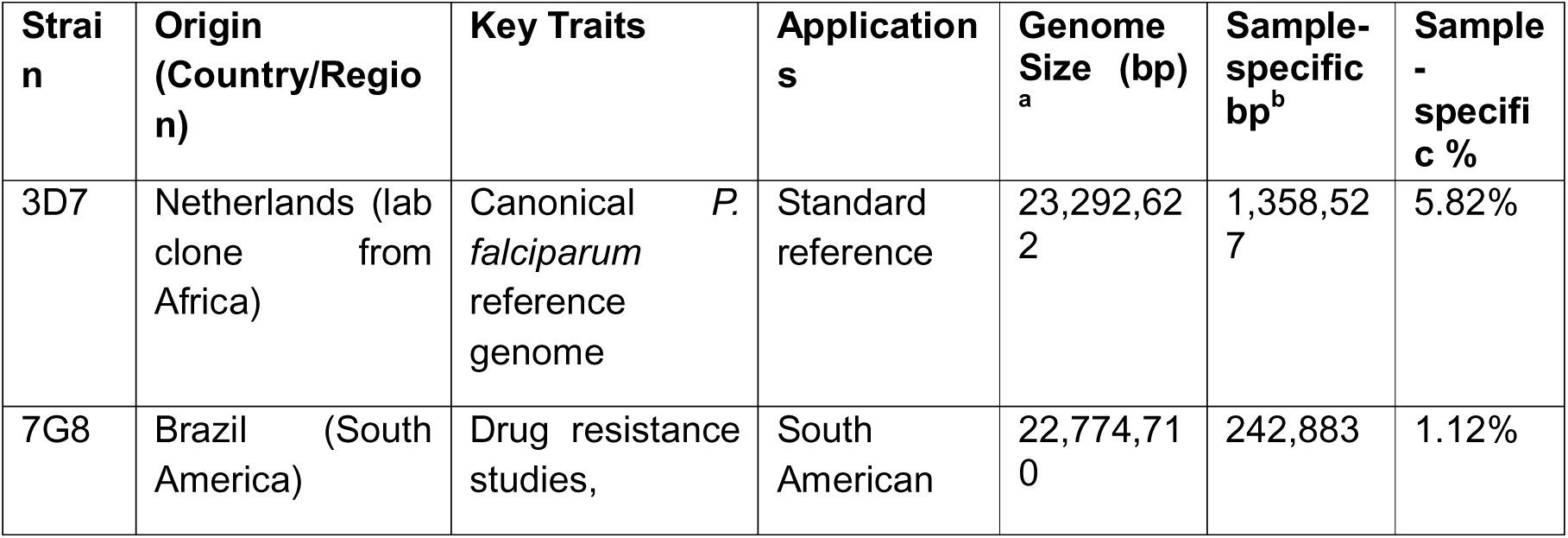
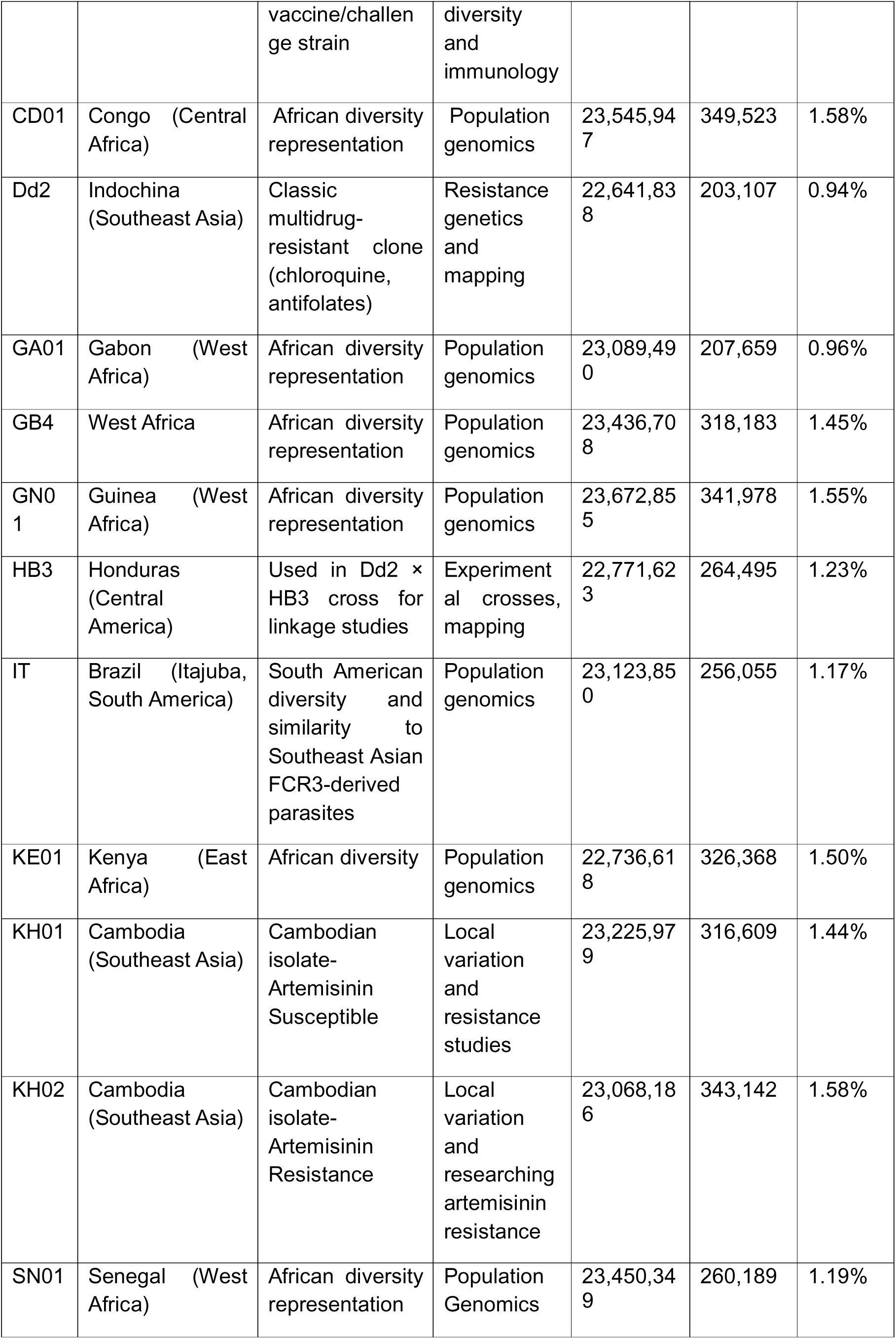
*P. falciparum* strain assemblies used to construct our pangenome (PfPan)

### Benchmarking Small and Structural Variant Calling

Next, we assessed the accuracy of small variant (<50 bp) and structural variant (SV) (≥50 bp) genotyping with PfPan relative to a conventional linear-reference mapping approach, using assembly-derived variant calls as the benchmark. Genotyping accuracy was evaluated using the F-measure, the harmonic mean of precision (the proportion of called variants that are true positives) and recall (the proportion of true variants detected). Values range from 0 to 1, with 1 indicating perfect concordance with the benchmark; the metric is undefined when there are no true positives (precision = recall = 0). Short reads from each sample were mapped either to the linear 3D7 reference using BWA or called from surjected alignments from PfPan, with variants called using GATK (small variants) or Delly, Manta and Dysgu (SVs). Variants were also compared to variants extracted from the assembly-based PfPan graph using VG Giraffe or genotyped directly from the graph (VG Call). To assess generalisability to samples not represented in the graph, we performed leave-one-out validation (LOOV), iteratively rebuilding PfPan with each sample excluded in turn and then mapping and genotyping the excluded sample’s reads against the resulting reduced graph^16–18^.

Pangenome-based approaches showed strong performance for both small variant and SV calling/genotyping across high-confidence genomic regions (see **Methods**). The Cactus pangenome graph, which genotypes variants directly from assemblies and therefore shares a representation with the assembly-derived truth set, achieved high F-measures for both SVs (median F-measure: 0.767; median recall: 0.792; median precision: 0.742) and small variants (median F-measure: 0.937; median recall: 0.966; median precision: 0.910) within the core genome (**Figure 1**). SV genotyping with VG Call outperformed conventional linear-reference-based methods for SV detection in the core genome, achieving a median F-measure of 0.512 (median recall: 0.595; median precision: 0.658), compared with 0.484 (recall: 0.337; precision: 0.870) for Manta, 0.353 (recall: 0.228; precision: 0.818) for Dysgu, and 0.107 (recall: 0.057; precision: 0.929) for Delly. VG Call achieved the highest F-measure and recall despite being limited to genotyping SVs represented in the pangenome graph, although its precision was lower than that of the conventional methods. Using the surjected alignments for *de novo* SV calling versus linear mapping led to increases in performance for both Delly (median F-measure: +0.104, recall: +0.062, precision: −0.032) and Dysgu (median F-measure: +0.053, recall: +0.060, precision: −0.100), with comparable performance for Manta (median F-measure: +0.020, recall: +0.029, precision: −0.059). In each case, the gain arose from improved recall at the cost of precision.

In contrast, linear mapping with GATK variant calling marginally outperformed surjection followed by GATK for small variants (median F-measure: 0.979 vs 0.972; median recall: 0.972 vs 0.970; median precision: 0.984 vs 0.974). The genotyping of small variants directly in the graph with VG Call performed less well (median F-measure: 0.759; median recall: 0.648; median precision: 0.924), reflecting a loss of sensitivity due to the genotyping approach rather than an excess of false positives.

Where performance was moderate, this may reflect differences in variant representation between callers rather than true errors. Investigation of discordant calls across the core genome and coding regions revealed that both GATK and VG genotyping calls had adequate median depth (GATK: 12–151x; VG Call: 8–31x) and quality scores (QUAL, a Phred-scaled estimate of the probability that the variant call is incorrect; median QUAL: GATK 188–2,537; VG Call 118–417), with median genotype quality scores (GQ, a Phred-scaled confidence in the assigned genotype) also high for VG calls in particular (median GQ: 71–256 vs. GATK median GQ: 75–99). Most discordant calls in both approaches had a variant allele fraction at or near 1.0 (VAF, the proportion of reads supporting the alternate allele; median VAF: 0.82–0.93), indicating these are high-confidence homozygous alternate calls rather than low-quality noise. Together with the high sequencing depth and quality metrics, these findings suggest that many discordant calls may represent genuine variants absent from the assembly-derived truth set, owing to missing loci or differences in variant representation between callers, rather than true sequencing or genotyping errors.

### Landscape of Genomic Variation Across PfPan (13 reference genomes)

After demonstrating the performance of PfPan for small variant and SV genotyping, we next explored genomic variation across the 13 reference genomes included in the pangenome. After removing variants called in only one of the 13 genomes (missing in 12 or more) and monomorphic sites (alternative allele frequency of 0), a total of 423,865 variants remained. This included 152,640 SNPs, 150,890 small insertions, 113,884 small deletions (absolute lengths < 50 bp), 3,859 structural insertions and 2,592 structural deletions (absolute lengths ≥ 50 bp) (**Figure 2A**). The median allele frequency was highest for SNPs (Median AF = 0.17) and small deletions (Median AF = 0.17), followed by structural deletions (Median AF = 0.09), small insertions (Median AF = 0.08) and structural insertions (Median AF = 0.08) (**Figure 2B**). A total of 236,124 variants were singletons and observed in only one of the 13 reference genomes. Variants were unevenly distributed across the *P. falciparum* genome. Most variants were observed in either coding regions (97%) or the core genome (88%) (**Supplementary Figures 3–5**). Notably, the core genome covers >85% of the total genome, making the variant density approximately 19 variants per kb. As expected, the variant density was highest for subtelomeric hypervariable regions (76 variants per kb) (**Supplementary Figure 6**).

**Figure 2.**
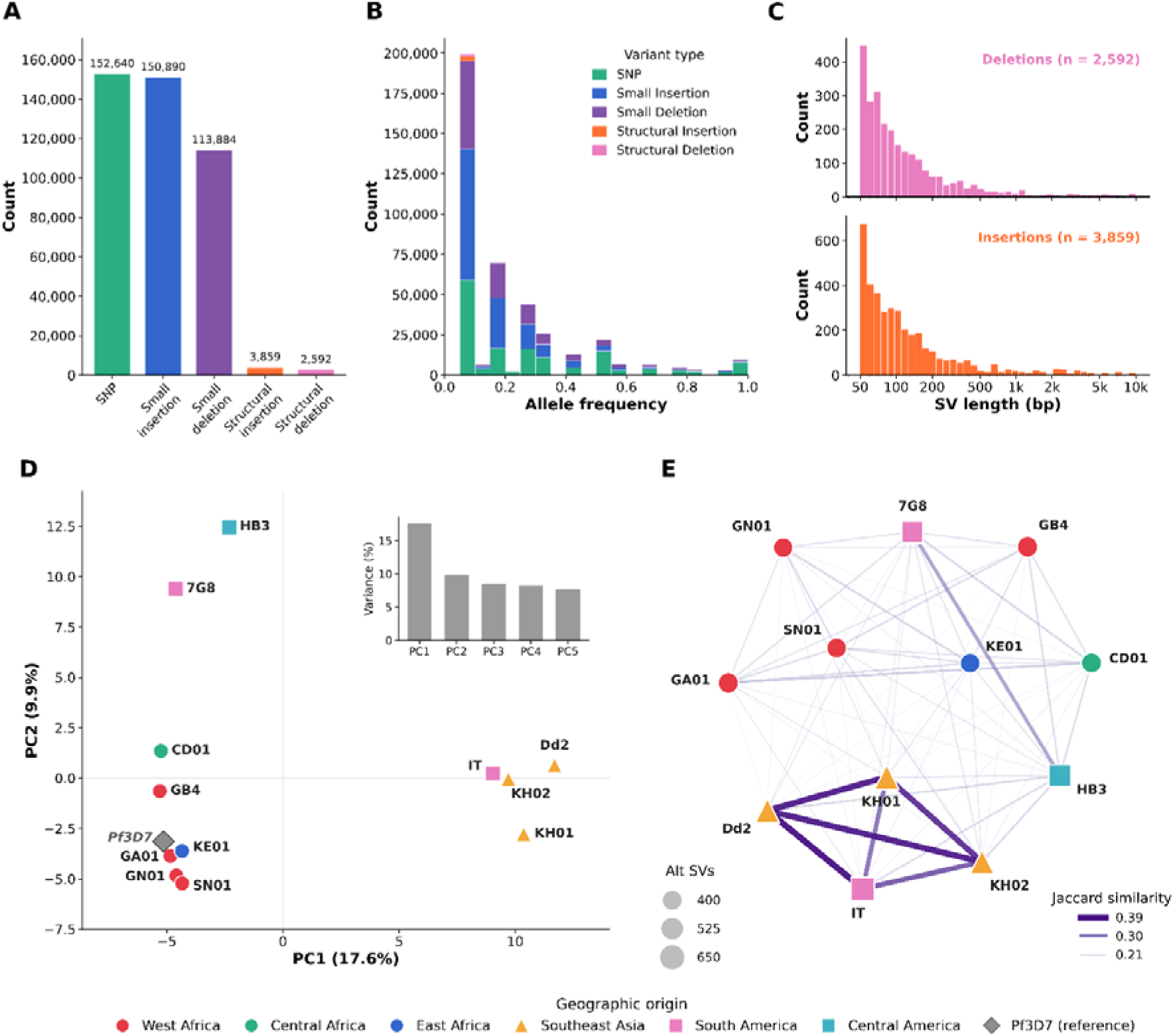
Genomic landscape of the *P. falciparum* pangenome (PfPan). **(A)** Bar chart showing the total number of SNPs, insertions, deletions, structural insertions, and structural deletions identified across the 13 PfPan reference strains. **(B)** Allele frequency distributions stratified by variant type, illustrating the site frequency spectrum across SNPs, insertions, deletions, structural insertions, and structural deletions. **(C)** Length distributions of structural insertions and deletions (≥50 bp). **(D)** Principal component (PC) analysis of the 13 reference strains based on a structural variant (SV) presence/absence distance matrix, with clustering reflecting the geographic origin of strains. **(E)** Network graph illustrating pairwise Jaccard similarity between reference strains based on shared SV haplotypes, with node size proportional to the total number of alternative allele SV calls per strain.

In all regions, SNPs and small insertions or deletions (indels) were the predominant variant types, whilst structural indels were rare (<5%). The internal hypervariable regions had the highest proportion of structural insertions (1.7%) and structural deletions (1.9%). In addition, the lengths of different variant types varied widely. Small insertions had a median length of 4 bp, while small deletions had a median length of 6 bp (**Figure 2B**). Structural insertions were expectedly larger, with a median length of 92 bp, and structural deletions had a median length of 88 bp (**Figure 2C**). Overall, these results reveal that genomic variation in PfPan is strongly influenced by genomic context, with the core genome and subtelomeric hypervariable regions exhibiting distinct patterns of variation in terms of variant type, size, and frequency. To explore the relationships among *P. falciparum* strains based on SV content, we performed principal component analysis (PCA) and constructed a Jaccard similarity network. SVs with absolute length ≥50 bp and present in ≥2 samples were retained (*n =* 1,586). The *P. falciparum* strains clustered according to their geographic origin, forming three main clusters: African, South American and Southeast Asian (**Figure 2D, Supplementary Figure 7**). This grouping was independently confirmed by the Jaccard similarity network, in which edge weights reflect the proportion of shared alternative calls between each strain pair (Jaccard index) (**Figure 2E**). Dd2, IT, KH01 and KH02 formed a distinct cluster with notably higher pairwise similarity (Jaccard Similarity: 0.34–0.39) compared to all other pairs (Jaccard Similarity: 0.20–0.26), consistent with a shared SV ancestry. Notably, IT originated from South America but can genetically resemble Southeast Asian FCR3-derived parasites due to early cross-contamination ^19,20^.

### Exploring Biologically Important Variation in PfPan (13 reference genomes)

We next sought to identify variants across 26 genes in PfPan (13 reference genomes) comprising both validated drug targets and candidate resistance loci (**Supplementary Table 1**). All strains demonstrated expected drug-resistant haplotypes at *pfcrt*, *pfdhfr*, *pfkelch13* and *pfdhps* (**Supplementary Figures 8–11**). Across all drug resistance loci, genetic variation was predominantly composed of small-scale mutations, including SNPs and small indels (**Supplementary Table 2**). The highest number of variants was observed in *pfubp1* (*PF3D7_0104300,* ubiquitin carboxyl-terminal hydrolase 1), which contained 80 SNPs, 47 small insertions, and 50 small deletions (reported per unique variant position). Other loci with substantial variation included *PF3D7_1025000* (62 SNPs, 37 small insertions, 40 small deletions), *pfmca2* (*PF3D7_1438400,* metacaspase-like protein; 55 SNPs, 33 small insertions, 23 small deletions), *pfcrt* (*PF3D7_0709000;* chloroquine resistance transporter; 33 SNPs, 25 small insertions, 21 small deletions), and *pfcoronin* (*PF3D7_1251200*; 39 SNPs, 23 small insertions, 22 small deletions). In contrast, SVs were rare. A total of 15 structural events were detected across 9 drug resistance loci, comprising 7 structural insertions and 8 structural deletions. Only 2 loci had multiple SVs, including *pfap2-mu* (*PF3D7_1218300,* AP-2 complex subunit mu), which contained both a structural insertion and a structural deletion at position Pf3D7_12_v3: 717118, and *pfubp1*, which contained both a structural insertion (Pf3D7_01_v3: 190952, Length: 60bp, Strain: KH02) and deletion (Pf3D7_01_v3: 187961, Length: 298 bp, Strains: Dd2 and 7G8).

All other affected loci contained only a single structural event each, including structural deletions in *pfdhfr* (*PF3D7_0417200,* DHFR-TS, bifunctional dihydrofolate reductase-thymidylate synthase; Pf3D7_04_v3: 747368, Length: 254 bp, Strains: GB4 and HB3), *pfdhodh* (*PF3D7_0603300,* dihydroorotate dehydrogenase; Pf3D7_06_v3: 132794, Length: 67–71 bp, Strains: CD01 and HB3), and *pfcrt* (Pf3D7_07_v3: 404179, Length: 52 bp, Strain: 7G8), as well as structural insertions in *pfcarl* (*PF3D7_0321900,* cyclic amine resistance locus protein; Pf3D7_03_v3: 922055, Length: 59 bp, Strains: GB4 and KH02) and *pfprs* (*PF3D7_1213800,* Pf3D7_12_v3: 589694, Lengths: 185–188 bp, Strains: 7G8, CD01, Dd2, GA01, GB4, GN01, IT, HB3, KE01, KH01, KH02 and SN01). In most cases, SVs were distributed across individual strains and are unlikely to play a role in drug resistance. However, the insertion at Pf3D7_01_v3: 190952 is of particular interest, as *pfubp1* has been hypothesised to be involved in resistance and KH02 is the only artemisinin-resistant strain included in PfPan. However, this variant was observed across artemisinin-susceptible samples in downstream analyses. The deletion at Pf3D7_07_v3: 404179 is also noteworthy given that 7G8 is chloroquine-resistant. Despite this, other chloroquine-resistant strains do not carry this deletion, suggesting it may not be generally important for resistance. Regardless, SVs remain an important source of genetic diversity, as they occur across multiple strain backgrounds and may contribute to the emergence of drug resistance or other adaptive phenotypes in specific contexts.

We further sought to identify SVs within complex regions of the *P. falciparum* genome. We identified ‘complex SVs’ characterised as clusters of multi-allelic SVs >1,000 bp in length (<10 kb) (see **Methods**). A total of 94 genomic windows containing 272 sites with 544 alleles were identified. Most complex SVs were observed in known hypervariable and/or multi-family genes, including *var*, *rif*, *emp3*, *mc-2tm*, *acs*, and *epf* family members (*epf1*–*epf4*), often occurring in loci containing multiple overlapping gene families (**Supplementary Table 3**). Intriguingly, we observed complex SVs across genes involved in gametocytogenesis. This included *pfgdv1* (*PF3D7_0935400*), *pfg27/25* (*PF3D7_1302100*) and *pf11-1* (*PF3D7_1038400*). We also observed complex variants in the GCH1 locus (*pfgch1*, *PF3D7_1223600*, GTP cyclohydrolase 1; Pf3D7_12_v3: 974,372–975,541), which contains multiple structural insertions, including a 2,216 bp (KH02) and an 861 bp (KE01) insertion at position 974,248, located upstream of *pfgch1.* Further analysis of *pfgch1* revealed additional large-scale SVs upstream of *pfgch1* at position Pf3D7_12_v3:968881 including in 7G8, Dd2 and GB4 (all >9 kb). Larger-scale complex loci were characterised further in the copy number variant analysis below.

We additionally searched for any SVs in the *pfhrp2* (*PF3D7_0831800*) and *pfhrp3* (*PF3D7_1372200*) genes, where deletions observed in Africa and South America are known to cause false-negative rapid diagnostic test (RDT) results (**Supplementary Table 4**) ^21–24^. No large-scale, gene-wide deletions were observed. However, we observed multiple variant types across the two genes. The *pfhrp3* gene contained 1 structural insertion, 2 small insertions, 2 structural deletions, 7 small deletions, and 33 SNPs, while *pfhrp2* contained 2 structural deletions, 3 structural insertions, 9 small insertions, 11 small deletions and 46 SNPs. Deletions upstream of the coding sequence for *pfhrp2* were observed in KH01, KH02, HB3, IT and GN01 (Range: 718–730 bp) (**Supplementary Figure 12–13**).

### Structural Variant Genotyping Across Diverse *P. falciparum* Populations

After mapping *P. falciparum* short reads to the pangenome, we first sought to demonstrate the utility of the pangenomic approach for identifying SVs that are common across and between populations. Given that pangenome-based SV genotyping outperformed linear *de novo* calling in the benchmarking analysis, SV calling was performed by genotyping against the pangenome graph rather than independent SV calling from short reads, meaning all variants reported here are also found in the pangenome assemblies. In total, post-filtering we identified 61,128 variants across 878 convenience samples from the public domain (Southeast Asia: 56.0%, West Africa: 21.3%, East Africa: 9.1%, Oceania: 4.3%, Central Africa: 4.1%, South America: 2.8%, South Asia: 2.3%). Of these, 82 were classified as structural deletions (median length 72 bp, range 51-423 bp) and 130 were classified as structural insertions (median length 67 bp, range 50-4,935 bp), post-filtering for SV length <10,000 bp.

To identify regional differences in structural variation, we investigated the top 1% of SVs by fixation index (Jost’s D), comparing each population to samples across all other populations **(Supplementary Table 5)**. We observed 6 SVs (3 deletions and 3 insertions), occurring across genes or intergenic regions. In Southeast Asia, high Jost’s D values were observed for a 198 bp insertion in the gene encoding transcription factor AP2-G (*pfap2-g*, 198 bp insertion at position Pf3D7_12_v3:914425). In addition, a 57bp insertion at position Pf3D7_03_v3: 222288 (Jost’s D = 0.63) in the *pfcsp* gene encoding circumsporozoite protein (CSP), as well as a 150bp insertion in *PF3D7_0620000* (Jost’s D = 0.80), demonstrated strong differentiation in West Africa. Other highly differentiated SVs included a 60 bp deletion in *PF3D7_1238700* in South Asia (Pf3D7_12_v3:1606570, Jost’s D = 0.26), a 63 bp deletion in *PF3D7_0935900* (REX1) in East Africa (Pf3D7_09_v3: 1420953, Jost’s D = 0.62), and a 60 bp deletion in *PF3D7_1417600* in Oceania (Pf3D7_14_v3:731919, Jost’s D = 0.65). These findings suggest that genes involved in mosquito-stage development and erythrocyte invasion may be subject to geographically variable selective pressures. None of the highly differentiated SVs overlaps with known drug resistance loci.

### Small Variant Discovery Across Diverse *P. falciparum* Populations

Although SVs were identified using a genotyping-based approach, the strong benchmarking performance of small-variant callers such as GATK enabled reliable small-variant discovery from the pangenome. This approach involved ’surjecting’ the graphical representation of the BAM file and undertaking a traditional ’linear’ variant calling approach. Across the 878 samples, 1,673,825 SNPs, 693,935 insertions and 696,633 deletions were called. Multidimensional scaling (MDS) of a SNP distance matrix demonstrated the expected geographic clustering, consistent with the 3D7-based linear mapping approach, with increased separation across the first principal component (**Figure 3**). Restricting the MDS to novel pangenome sequence recovered the same broad population structure seen using the full variant set (**Figure 3**). This indicates that novel sequence content captured by PfPan contributes to fine-scale population structure. We next performed a small variant-based population differentiation analysis, using Jost’s D to identify SNPs and indels exhibiting strong allele-frequency differentiation between geographic populations. A total of 10,674 SNPs, 9,665 deletions and 6,744 insertions were amongst the top 1% of fixation indices across all geographic regions. A combined total of 49,342 sites met the fixation thresholds, comprising 28,170 SNPs, 14,187 insertions and 6,985 deletions.

**Figure 3.**
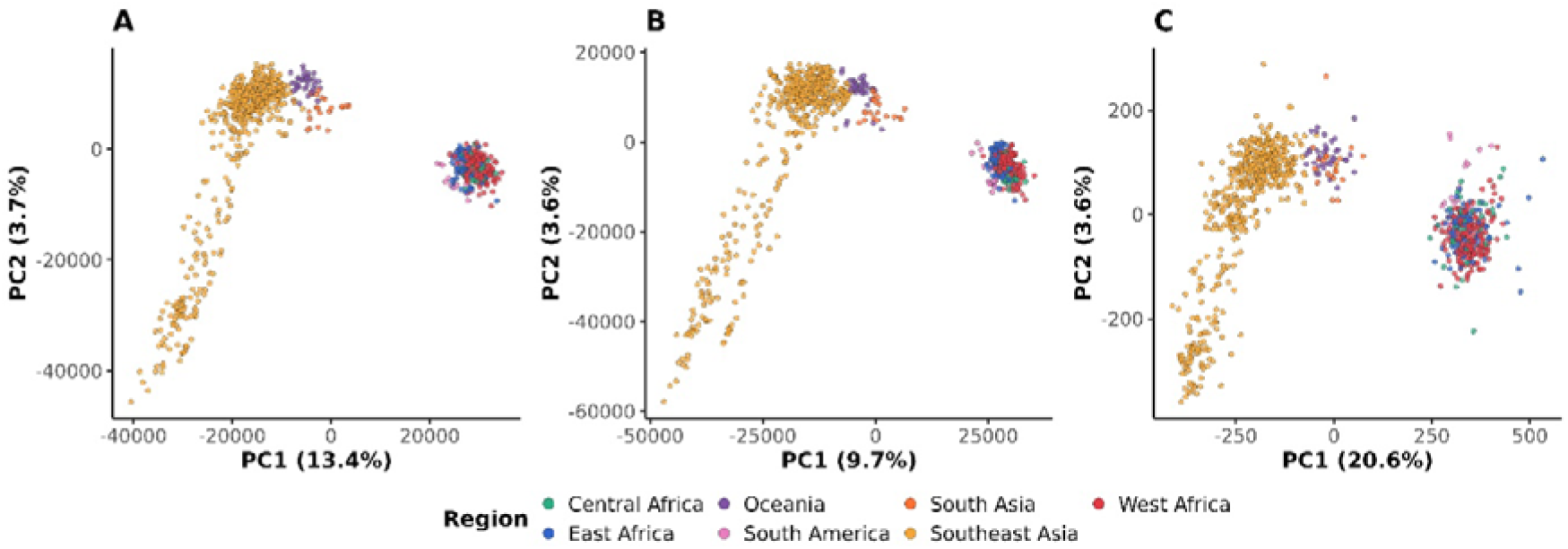
Population structure of 878 global *P. falciparum* isolates based on biallelic SNPs called from the PfPan pangenome and linear reference mapping. Multidimensional scaling of pairwise genetic distances for *n* = 878 isolates, based on **(A)** all variants called against the pangenome graph using GATK, **(B)** variants called against the linear reference using GATK, and **(C)** corroborated novel insertions only. Each point is one isolate, coloured by geographic region of origin. Axes show the first two principal coordinates, with the percentage of total variance explained.

Given that F_ST_-based analyses of SNPs have been extensively reported in the literature, we focused on indels to complement existing knowledge and further characterise small SV calls (**Supplementary Table 6**) ^25^. From this analysis, we observed 17 indels in *pfcrt* (CRT) ranging from -16 bp deletions to 14 bp insertions predominantly in Oceania, African regions and South America, with the strongest differentiation observed for a 9 bp deletion at Pf3D7_07_v3:405673 in Oceania (Jost’s D = 0.95; alternative allele frequency 95.7% vs 0.1% globally). In addition, two variants in the coding and intergenic regions of *pfkelch13* (K13) were observed, including a 6 bp coding insertion (Pf3D7_13_v3:1726571 G>GATTATT) at 0.61 allele frequency in Southeast Asia and a 3 bp upstream variant (Pf3D7_13_v3:1728023 T>TATA) at 0.59 allele frequency in South America. These variants were not unique to either region or were also present in isolates carrying artemisinin-susceptible genotypes, suggesting they are unlikely to represent primary resistance determinants. The intergenic regions surrounding the *pfdhfr* locus (DHFR-TS) also contained indels, including 14–16 bp repeat-length variants observed in Oceania and South America. These occurred alongside an 8 bp insertion and a 6 bp deletion in Oceania, with all variants occurring in intergenic AT-rich sequence and demonstrating elevated differentiation (Jost’s D = 0.24–0.92).

Additional differentiated variants were identified in *pfap2-mu* (AP2-MU), where a 3 bp deletion was observed in 83.8% of samples from Southeast Asia (Jost’s D = 0.71), and in *pfubp1* (UBP1), where a 9 bp deletion was present in 73.7% of samples from Oceania (Jost’s D = 0.61). Variants were also detected in *pfmdr1* (MDR1), *pfpi4k* (PI4K), and *pfmca2* (MCA2), including deletions of up to 24 bp that were most frequent in South America and Southeast Asia, further supporting the presence of geographically structured indels across multiple loci with potential links to drug resistance (**Supplementary Table 7**).

### Copy Number Variation

Detecting copy number variants (CNVs) from pangenome-derived alignments depends on how the duplication is represented in the underlying assemblies, and the two cases require different approaches. Where a duplicated unit is collapsed into a single copy during assembly, it is not represented as a SV in the graph; reads from every copy in a sequenced isolate align to that one position, and copy number remains recoverable from coverage. Where the unit is resolved, the copies are represented as distinct traversals within the graph and must be characterised from the graph’s own allele representation. To demonstrate the first case, we focused on *pfmdr1* amplifications, which are well characterised in *P. falciparum* given their role in drug resistance ^4^. Using short-read data from the Pf8 dataset (N=878), we took a targeted coverage-based approach across established *pfmdr1* breakpoints and compared pangenome-derived calls against those from the linear mapping pipeline ^4^. High-confidence pangenome-based calls showed high concordance with linear mapping (overall concordance = 0.997; sensitivity = 96.2%; specificity = 100%; **Supplementary Table 8**), with no false-positive pangenome-only calls. The two missed amplifications had borderline coverage ratios near the calling threshold. Short-read mapping to a pangenome graph is therefore a viable and accurate alternative to linear reference-based approaches for detecting clinically relevant CNVs of this class. However, using a coverage-based approach, amplifications in the drug-resistance gene *pfgch1,* which is known to have multiple copies across some strains, were not detectable ^15^. Here, the reads are consistent with both reference and alternative paths and are therefore assigned arbitrarily with mapping quality near zero (**Supplementary Figure 14**), so the signal does not survive surjection of the graph alignments onto linear coordinates.

Therefore, we characterised *pfgch1* from the founder assemblies and the graph’s allele representation instead (**Supplementary Table 9**). To characterise these variants, we extracted inserted and deleted sequences >500 bp in length and BLASTed them against the 3D7 reference to capture CNVs that depth-based approaches may miss. Using BLAST characterisation, a 10,418 bp insertion at position 968,881 (Pf3D7_12_v3) was identified spanning *PF3D7_1223800, PF3D7_1223900* and *pfgch1* (*PF3D7_1224000*), with BLAST alignment to the reference genome yielding 98.5% identity across 100% query coverage (Dd2-like). The insertion adds two further copies of a ∼5.2 kb unit, giving three copies in total, of which the middle copy is in inverted orientation (**Figure 4**). A second, larger insertion of 14,505 bp (two extra copies of a larger 7.3 kb unit) at the same anchor position spans the same three genes with 98.4% identity, as well as approximately 85% of *PF3D7_1223700*, suggesting a further expansion of the duplicated unit (GB4-like). A third, insertion of 2,216 bp at position 974,248 maps within *pfgch1* and its flanking sequence (98.8% identity; 100% query coverage), representing a distinct SV (KH02-like). A fourth, smaller insertion of 861 bp at the same position lies immediately upstream of *pfgch1* (99.8% identity; 100% query coverage) and comprises two tandem copies of a ∼440 bp unit (KE01-like). A larger duplication at this locus in 7G8 was not analysed, as its path through the graph is interrupted within the array. To confirm they reflect genuine duplications rather than artefacts of graph construction, we mapped short-read data for the founder strains linearly to the 3D7 reference and examined read depth across the locus. Coverage was elevated in precisely those strains carrying an insertion in the graph and flat at single copy in the remaining sequences (**Supplementary Figure 15**).

**Figure 4.**
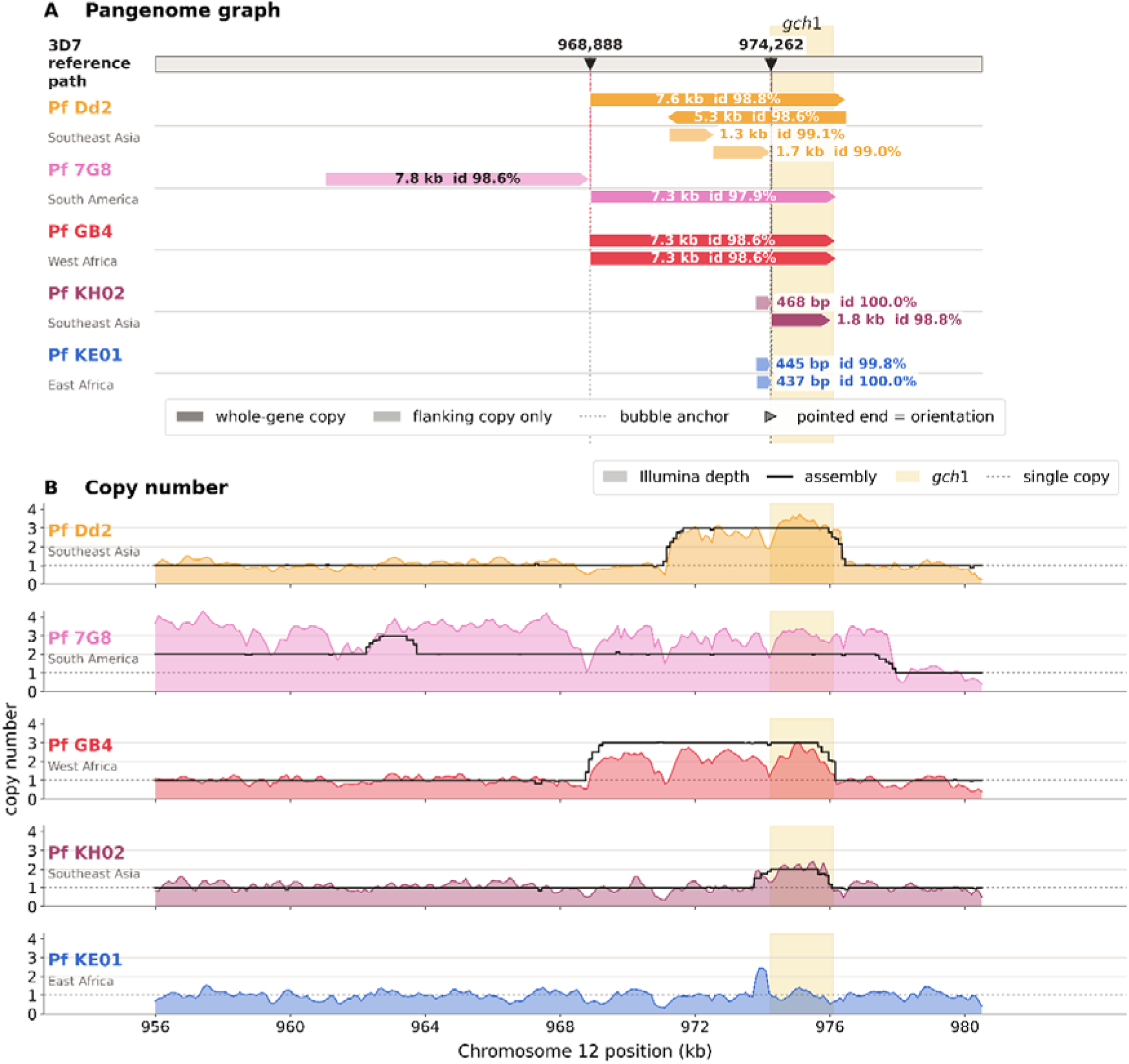
Complex structural variation surrounding *pfgch1* reveals large-scale gene duplication across global *P. falciparum* populations. (**A**) The gene track of the 3D7 reference covering *pfgch1*. The dotted vertical lines mark the two bubble anchors at which non-reference paths diverge from and rejoin it (Pf3D7_12_v3:968,888 and 974,262). For each assembly carrying additional sequence, that sequence was realigned to the reference window, and each resulting block is drawn at the reference interval. Pointed ends indicate orientation; solid fill denotes a copy spanning ≥90% of *gch1*, pale fill a copy of flanking sequence only. **(B)** Copy number across the same interval. Filled areas show Illumina read depth in 100 bp bins, normalised to the median depth of the local single-copy flanks (915,000–945,000 and 980,000–1,020,000 bp). The solid black lines show copy number derived independently from the PacBio assemblies. The dashed line marks single copy number, and the shaded band highlights *pfgch1* (974,225–976,097 bp).

Beyond *pfgch1*, this analysis identified recurrent CNVs at several biologically significant loci that also overlapped with sites enriched in SV signal noted above (‘complex loci’) (**Supplementary Table 10**). Insertions were identified at *pflsa1* (*PF3D7_1036400*) across multiple samples, suggesting high-level tandem amplification of this liver stage antigen. The deletions in *pfgdv1* (*PF3D7_0935400*; 21,232 bp and 5,862 bp) were located within the coding sequence, implicating structural variation in the regulation of sexual commitment and transmission. Additional notable deletions were identified encompassing *pfclag3.1* and *pfclag3.2*, *pfrh2b*, members of the EPF exported protein family, as well as deletions in the promoter region of the gene encoding KAHRP, which have been described previously ^6^. Whilst these findings demonstrate that analysing large insertions and duplications (>500 bp) can improve the characterisation of copy number variation within complex genomic loci, calling such variation in lower coverage field samples using the pangenome remains challenging.

## DISCUSSION

Here, we present PfPan, a high-quality *P. falciparum* pangenome comprising 13 geographically diverse reference genomes, including the widely used 3D7 strain. We demonstrate the improved ability of pangenomes for SV genotyping and detection, and comparable *de novo* calling of small variants through comparisons with linear variant calling methods. High concordance was observed between variants called in the core genome across high confidence callsets. We note that genotypic discordance or lower performance metrics could be attributable to the differing qualities of the underlying reference genomes. Not all assembled reference genomes had chromosomes with telomeres attached, and there were variations in the number of *var* genes reported ^6^. Even so, combining these reference genomes together enables us to overcome a fundamental flaw of genome analysis: reference bias. 3D7 is a robust reference genome for *P. falciparum*, particularly for core genome-based analyses, and has been widely used across population genomic studies ^25^. However, the addition of 4.77 Mb of novel sequence, together with a graph-based genome representation, enables the discovery of more complex features of the *P. falciparum* genome that would likely be missed when relying on a single reference alone.

A key advantage of pangenome approaches is the ability to identify and genotype SVs, which have historically been difficult to resolve due to reliance on short-read sequencing data and single-reference genomes. Although *de novo* calling SVs was possible using tools such as Manta and Dysgu, we showed that genotyping SVs within the pangenome graph performed on par or marginally better than traditional mapping and variant calling approaches. In this study, we identified more than 5,000 SVs (≥50 bp) across the 13 reference genomes in PfPan. Several of these SVs have important biological relevance, with existing literature supporting their functional roles. We focused particularly on “complex loci”, regions of the genome enriched for SVs that have historically been challenging to analyse. As expected, complex loci included hypervariable gene families such as *var, rif, emp3, mc-2tm, acs* and *epf* family members (*epf1–epf4*). The inclusion of multiple references in the pangenome accounts for known SVs during read mapping, reducing false SNP calls that arise from structural misalignments to a single reference ^5^. Genes involved in gametocytogenesis were also amongst complex loci, including *pfgdv1, pfg27/g25* and *pf11-1.* GDV1 is a regulator of sexual commitment that induces gametocyte development by antagonising HP1-dependent gene silencing, and large deletions (18*–*26 kb) encompassing the *pfgdv1* locus have been associated with complete loss of gametocyte production in laboratory strains ^26^. *Pfg27/25* encodes an abundant gametocyte-specific protein and exhibits upstream deletions in laboratory lines in addition to structural polymorphisms in repetitive sequences that affect gene regulation. Despite this, prior studies indicate *pfg27* knockout parasites can still complete gametocytogenesis ^27^. *Pf11-1* is involved in erythrocyte rupture during gametogenesis and frequently undergoes chromosome breakage and healing events that delete >90% of the gene in laboratory strains through telomere addition ^28^. The identification of SVs within these critical gametocytogenesis pathways further validates the pangenome approach as a powerful framework for biologically meaningful structural variation discovery.

We also report SVs with potential clinical and biological relevance. For example, a 254 bp deletion upstream of drug-resistance gene *pfdhfr* was identified in both GB4 and HB3. HB3 carries the *pfdhfr* S108N mutation, a validated marker of pyrimethamine resistance, whereas GB4 shares the same *pfdhfr* coding-region haplotype as 3D7 ^30^. This suggests that the 254 bp deletion is not restricted to drug-resistant genetic backgrounds, although it was absent from all 878 field samples. Other SVs demonstrated differentiation at the regional level, including deletions in *pfrex1*, *PF3D7_1417600* (ookinete surface and oocyst capsule protein) and *PF3D7_1238700* (BTB/POZ domain-containing protein, putative), and insertions in *pfcsp, PF3D7_0620000* (PIMMS43) *and pfap2-g.* Whilst some of these genes are implicated in host cell remodelling and invasion (*pfrex1* and *pfcsp*), highly differentiated variants in other genes may represent regional adaptation to mosquito vectors and transmission ^31–33^.

We next sought to investigate CNVs in drug-resistance genes using targeted coverage-based approaches based on validated breakpoints and BLASTing large-scale SVs. CNV calls at *pfmdr1* were largely consistent with previous reports ^34,35^. The *pfmdr1* copies are collapsed into a single copy, meaning that coverage-based approaches are still suitable for CNV detection in non-complex loci ^6^. In contrast, *pfgch1* and its upstream regions were identified among complex loci enriched for SVs >1 kb in PfPan, capturing previously described copy number variation at this locus. ^15^. The additional copies/ tandem repeats are not collapsed and are instead represented as alternative paths across the *pfgch1*. Therefore, when mapping to the pangenome at highly similar non-collapsed duplicated sites, reads may map well to multiple paths, reducing the overall mapping quality due to ambiguity. This observation is reflected in downstream surjection of pangenome-based alignments to a single linear reference. Consequently, conventional coverage-based CNV estimates may not fully distinguish a true amplification from an inserted homologous sequence.

To characterise smaller duplications at ‘complex loci’, we BLASTed indel sequences (>500 bp). This identified the high-frequency 10,418 bp insertion as having three gene hits: *PF3D7_1223800, PF3D7_1223900* and *pfgch1*. This observation has been reported previously using long-read sequencing and was associated with *pfdhodh* copy number ^37^. We also report deletions upstream of the coding sequence for *pfhrp2*. Deletion of *pfhrp2* has been associated with false-negative RDT results and represents a significant diagnostic evasion mechanism with direct public health implications ^22–24^. The inclusion of new assemblies from South America and the Horn of Africa in PfPan in the future would therefore be essential to effectively genotype large-scale variation at this locus. In addition, new methods to call CNVs encoded as structurally complex loci in pangenomes would enable further characterisation of these variants in field samples. Future analyses incorporating assembly-resolved breakpoints, long-read validation, and reference-free k-mer approaches will also be important to resolve the underlying architecture and CNV calling at these regions ^36^.

Moreover, several avenues for future research could further enhance the value of PfPan and our broader pangenome framework. First, we note that the use of a convenience sampling design for short-read mapping may underestimate the true global distribution of genomic variation. Monoclonal samples from Southeast Asia are the most numerous in the Pf8 dataset, so the eligible samples (monoclonal, non-sWGA) came disproportionately from this region, and sampling was not balanced across regions. Comparisons of allele frequency and genetic differentiation between regions therefore rest on unequal sample sizes and should be treated as exploratory. Future use of more balanced datasets with greater regional representation would allow fairer comparison of allele frequencies and genetic differentiation across regions. In addition, the set of high-quality reference strains incorporated into the graph does not yet fully capture the genomic diversity of *P. falciparum*.

As with previous pangenome studies, our primary objective was to develop a resource that reduces reference bias; however, incorporating additional *P. falciparum* assemblies in future iterations, particularly strains carrying clinically important variants such as *pfhrp2/3* deletions, will further advance this goal ^22,24^. Further, reducing the computational burden of mapping to a pangenome and building more robust pipelines for non-human species is essential. This will allow for more large-scale, holistic population genomic analysis in the future. Even so, several challenges remain. A large proportion of public WGS data for *P. falciparum* is short-read and has been made possible due to selective whole-genome amplification to increase the coverage of parasite DNA, making it challenging to assemble due to low or variable coverage. This difficulty could partially be resolved using long-read sequencing data for further SV discovery. We also note that the pangenome approach has only been applied to monoclonal field samples. Further benchmarking and the potential application of deconvolution methods should be carried out for polyclonal samples in the future ^38^. Here, we also use GATK for small variant calling as it is standard practice for *P. falciparum* genomic analysis. Investigating other variant callers such as DeepVariant in combination with pangenome mapping approaches could improve precision and recall ^8,39^. However, DeepVariant is pre-trained on human data and will require optimisation before being applied to *P. falciparum* genomes.

Overall, we present PfPan, the first graph-based pangenome reference for *P. falciparum.* We highlight that including additional reference genomes in addition to 3D7 as part of a pangenome can improve the accuracy of SV genotyping, refine CNV calls, and lead to comparable small variant discovery compared to linear reference genomes. These findings provide valuable insights into parasite structural variation and may inform future research into malaria drug resistance, genomic surveillance, and the identification of potential therapeutic targets, thereby contributing to improved strategies for controlling this high-burden disease.

## METHODS

### Pangenome Construction and Quality Assessment

To construct the *P. falciparum* pangenome (PfPan), we obtained 15 reference genome assemblies made available as part of the MalariaGEN Pf3k project and the standard *P. falciparum* 3D7 reference genome (version 3) ^6^. The 15 additional genome assemblies included 7G8, CD01, Dd2, GA01, GB4, GN01, HB3, IT, KE01, KH01, KH02, ML01, SN01, TG01 and SD01 strains. These assemblies were initially included in the first iteration of PfPan. Following inspection of alignment statistics, SD01 was excluded because of inconsistencies in contig size, while ML01 and TG01 were excluded after identifying evidence that these assemblies were derived from polyclonal (mixed infection) samples. PfPan was subsequently reconstructed without these assemblies. The final version of PfPan was constructed using the cactus-pangenome pipeline available as part of the Minigraph-Cactus toolkit (v2.9.3), using the 3D7 reference genome as the backbone genome^40^. In brief, this pipeline uses Cactus software to produce a hierarchical alignment (HAL) of all input assemblies. From this alignment, whole-genome and per-chromosome graph representations are generated in multiple graph formats (VG, GBZ, ODGI, and GFA) to use for visualisation or downstream processing. As part of the standard Minigraph-Cactus pipeline, an allele-frequency-filtered graph was generated using --filter 2, retaining only nodes and edges supported by at least two haplotypes. This filtered graph was generated as a standard pipeline output but was not used for downstream analyses. For the human pangenome, this produced a more compact graph optimised for read mapping and downstream analysis. However, as *P. falciparum* is relatively conserved and only 13 reference genomes were included in the pangenome, we opted to use a clipped graph, in which regions not incorporated into the underlying graph, such as unaligned flanking sequences, are removed or truncated. This retains most of the variation while simplifying the graph structure for visualisation and mapping purposes. The pipeline also produced a full graph, comprised of the complete set of sequences from all input assemblies, including unaligned centromeric or repetitive regions. This resulted in increased graph size and complexity, and consequently, was not used in downstream analyses. Additional outputs included variant call format (VCF) files anchored to the 3D7 reference genome and ODGI (optimized dynamic genome/graph implementation) visualisations (--viz) (see **Exploring Pangenome Variation**).

We used ‘vg stats’ to count the number of nodes, edges and paths within the PfPan graph (vg v1.61.0) ^18^. This was followed by an application of the Panacus toolkit to assess pangenome size and growth ^41^. We focused on countable graph features including nodes, edges and base pairs. The coverage of each feature was defined as the number of distinct paths (genomes) in which a given feature was present, and the quorum parameter was used to determine the proportion of paths required for a feature to be considered core (-l 1,2,1,1,1 and -q 0,0,1,0.5,0.1). The average node degree was calculated using ‘odgi degree’ (odgi v0.9.3) ^42^. Core and accessory genome fractions were estimated based on coverage and quorum thresholds. Pangenome growth curves were used to visualise how the number of common and core bases changed with the addition of samples using a modified version of the Panacus visualisation tool.

### Benchmarking

To benchmark the performance of the pangenome-based approach for small variant and structural variant (SV) discovery, we compared the pangenome-derived variant calls to assembly-based baselines and additionally evaluated results from a conventional linear reference-mapped approach. The assemblies of 7G8, CD01, Dd2, GA01, GB4, GN01, HB3, IT, KE01, KH01, KH02, and SN01 strains were mapped to the 3D7 reference genome using minimap2 (v2.28) and small variants (<50 bp) were called using paftools (v2.30) ^43^. In addition, SVIM-asm (v1.0.3, haploid) was used to detect SVs (≥ 50bp) from the assembly alignments ^44^. These callsets were used as a baseline for benchmarking performance. Three additional variant callsets were generated for benchmarking: (i) pangenome-derived variants generated as part of the cactus-pangenome pipeline; (ii) variants called from the pangenome after mapping short reads from the same strains to PfPan using a leave-one-out-validation (LOOV) approach (see **Mapping and Variant Calling from a Pangenome)**; and (iii) variants called using a linear mapping approach (see **Linear Mapping and Variant Calling**). To test the generalisability of the pangenome, we used a LOOV approach. The pangenome was reconstructed, omitting each validation sample in turn from the set of assemblies used to build PfPan, generating a strain-specific pangenome graph that excluded that sample’s own genome. Short reads from the omitted strain were then mapped to this reduced graph and variant genotyping/calling was performed, allowing us to assess genotyping performance for a sample not represented in graph construction.

Small variants (<50 bp) generated from linear mapping and pangenome approaches used 3D7 as the backbone reference. GATK (v4.1.4.1) callsets were separated into SNPs and indels with SelectVariants, and only ‘FILTER=PASS’ variants in the VCF files were retained; SNPs and indels <50 bp were merged and normalised ^16^. Variant quality score recalibration was not applied here to prevent bias in performance estimates, due to known high-quality existing SNPs. Alternative genotypes were assigned if ≥80% of the reads supported the alternative allele. Pangenome-derived SNPs were filtered to PASS, short indels <50 bp FILTER=PASS were separated, and the two sets were concatenated to produce the final small variant set; all callsets were further restricted to DP > 5 and homozygous alternate genotypes (GT = 1/1) to match the haploid paftools truth sets. Previous pangenomes have evaluated small variant discovery using high-confidence regions ^8^. To define a set of high-confidence regions, we performed (i) boundary trimming to remove contig-break artefacts, (ii) excluded high-variant-density windows (hypervariable regions), (iii) excluded known hypervariable gene families (*var*/*rif*/*stevor*) and centromeres, and (iv) excluded tandem-repeat regions that were found using both a genome-wide scan (Tandem Repeats Finder v4.10.0) and per-variant repeat-context annotation (vt annotate_indels v0.57721). We also focused on the core genome and coding regions. All VCFs were decomposed using RTG Tools (v3.13) (https://github.com/RealTimeGenomics/rtg-tools) (vcfdecompose, --break-mnps --break-indels) and benchmarked per strain with RTG vcfeval in annotate mode using - -all-records, --ref-overlap, --no-roc.

SVs (≥50 bp) were derived from Delly (v1.5.0) and pangenome callsets using 3D7 as the reference ^17^. Delly variants were restricted to records with a defined SVTYPE, and absolute SVLEN ≥50 bp, while pangenome-derived callsets were normalised and restricted to variants with FILTER=PASS and SVLEN ≥50 bp using bcftools (v1.12) ^45^. Prior to benchmarking, both baseline (SVIM-asm truth) and comparison callsets were restricted to ALT genotypes while retaining mixed genotypes. Benchmarking was performed per strain using Truvari (v5.3.0) with parameters: reference distance 1000 bp (-r 1000), maximum comparison distance 1000 bp (-C 1000), minimum size 50 bp (-s 50), maximum size 10,000 bp (--sizemax 10000), percent similarity threshold 0.3 (-P 0.3), minimum reciprocal overlap 0.0 (-O 0.0), and minimum sequence similarity 0.0 (-p 0.0) using the same aforementioned BED files for comparisons across genomic regions ^46^. The Truvari parameters were selected to tolerate breakpoint and representation differences in repetitive sequence; sensitivity to these settings is shown in **Supplementary Figure 16**.

### Exploring Pangenome Variation in PfPan (13 reference genomes)

We used the VCF file generated by the cactus-pangenome pipeline to explore variation within the pangenome. To produce the VCF file, the pangenome graph is decomposed into nested subgraphs (“snarls”) that each represent individual or multiple genetic variants, which are then transformed into a VCF output using vg deconstruct. We chose to use 3D7 (v3) as the backbone for the pangenome VCF, as this has the most comprehensive annotation available. The resulting VCF file was filtered to obtain high-quality variants that represent variation across the 13 *P. falciparum* assemblies included in the graph. Unlike traditional VCF files obtained through mapping approaches, this output does not contain any associated mapping or variant call quality metrics, so variants were filtered pragmatically according to methods previously employed by other pangenome studies ^12,40^. Multi-allelic sites were first normalised using bcftools (v1.12), before undergoing annotation with Truvari (v5.3.0) (anno svinfo) to add SV information, including type and length ^45,46^. We subsequently removed fixed or monomorphic sites, keeping only polymorphic variants. SVs >10 kb in length were excluded from the pangenome VCF analysis to avoid uncertain calls; larger events were instead characterised using BLAST-based approaches described below. The corresponding gene and predicted effect for each variant were annotated using bcftools csq to aid biological interpretation. All variants underwent additional manual classification, where variants <50 bp were labelled as ‘small variants’ and variants ≥50 bp in length were reported as SVs. We also characterised ‘complex SVs’, which were defined as genomic regions where large SVs (>1 kb) clustered within 200 bp windows, contained at least two alleles, and spanned a minimum of 1 kb, with overlapping genes identified using a 500 bp buffer.

To identify small variants and SVs within medically or biologically important regions, we filtered the VCF file for variants in drug resistance regions, vaccine candidates and other genomic regions of interest. These regions capture mutations across validated drug resistance gene candidates and suggested host-pathogen interaction candidates, as listed in the Malaria-Profiler tool and World Health Organization catalogues of drug resistance mutations ^47–49^. Variants of interest were prioritised based on their size, geographic distribution, population frequency, and potential association with relevant phenotypes. For example, we prioritised variants that were restricted to reference genomes originating from the same geographic region or that occurred in genomes with known drug resistance phenotypes, including partial artemisinin resistance. Variants of interest were visualised using a combination of methods, including Sequence Tube Map, odgi (v0.9.3) and Bandage software ^42,50,51^. To aid visualisation in Sequence Tube Map and Bandage, “vg” and “gfa” input files were obtained for each chromosome using ‘vg chunk’ and ‘odgi view’ respectively^18^. The region containing the SV was extracted using ‘vg chunk’. Variants of interest were confirmed using manual inspection of multi-sequence alignments between the 13 reference genomes obtained using minimap2 (v2.28) ^43^.

### Mapping and Variant Calling from a Pangenome

To demonstrate the advantages of a pangenome approach, we mapped global *P. falciparum* short-read (Illumina) data to PfPan. As selective whole genome amplification (sWGA) can lead to variable coverage, which impacts the detection of SVs, we selected samples that had undergone whole blood filtration. We also only included samples considered to be monoclonal. This was determined using 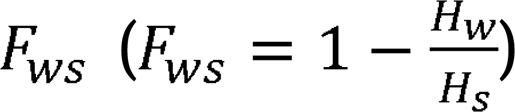, where *Hw* is the heterozygosity within a single infection and is the heterozygosity within a *P. falciparum* population ^52^. A threshold of 0.95 was used to indicate a monoclonal infection. This statistic was measured using the linear mapping and variant calling methods described below. All samples were sourced from the public MalariaGEN Pf8 dataset, selecting for samples with adequate coverage (at least 60% of the genome with coverage >5x, **Supplementary Table 11**). Due to the high computational demands for mapping and variant calling, we used a representative dataset, whereby the data underwent random stratified sampling to select a representative proportion of samples that met the criteria ^53^. We systematically selected 878 isolates by random sampling without replacement from 6,503 publicly available non-sWGA *P. falciparum* whole-genome sequences passing quality thresholds for minimum depth, percentage mapped reads, and clonality (F_WS_ > 0.95), retaining approximately 10% of samples. The overrepresentation of Southeast Asian isolates in the final dataset (56.0%), relative to their proportion in the pre-filtered pool (∼40%), likely reflects the well-established higher clonality of parasite populations in this low-transmission region, resulting in a greater proportion of Southeast Asian samples passing the F_WS_ threshold. The remaining isolates comprised samples from West Africa (21.3%), East Africa (9.1%), Oceania (4.3%), Central Africa (4.1%), South America (2.8%), and South Asia (2.3%). This sampling strategy was designed to support comparisons of SV genotyping and variant calling between pangenome-based and conventional linear-reference mapping approaches. In total, 878 samples were included in the analysis.

Prior to mapping, all raw reads underwent standard quality checks and filtering. Raw reads were trimmed using fastp (v1.0.1) and host reads were removed using Hostile (v2.0.0) ^54,55^. Short reads were mapped to PfPan using vg giraffe (vg v1.61.0) to output an alignment in graph alignment format (gaf) ^18^. After aligning to the pangenome, two forms of variant discovery were considered. Firstly, for SV genotyping from the pangenome directly, a coverage index was created using vg pack (-Q 5), followed by vg call specifying ‘3D7’ as the reference backbone and a ploidy of 1. To create a SV genotype callset (N=878) from PfPan, the resulting field sample VCFs were merged and normalised using bcftools (v1.12) and annotated using Truvari (v5.3.0) (anno svinfo) ^45,46^. Bcftools was also used to filter the multi-sample VCF for high-quality variants. Genotypes with ‘FMT/DP<5’ were encoded as missing. Sites were then retained if they had a quality score of at least 30, no more than 20% missing genotypes after encoding missingness, and a non-reference allele count of at least two, excluding singletons; variants ≥10 kb in length were discarded, as events of this size are not reliably genotyped from short-read data. SVs were retained if they were greater than or equal to 50 bp.

For *de novo* variant discovery, the mapped reads from the graph (gaf) were projected to a linear reference BAM format using vg surject (--interleaved and -R to assign read groups) and 3D7 paths. Here, 3D7 was chosen as the backbone reference due to its high assembly quality and annotation. The subsequent BAM file was subject to linear variant calling using GATK (v4.1.4.1) HaplotypeCaller (-ERC GVCF) and SV discovery using Delly (v1.5.0), Manta (v1.6.0) and Dysgu (v1.9.0) ^16,17^. Delly was run in two rounds, where the first was executed on individual samples to obtain per-sample calls. These calls were combined using Delly merge to create a BCF file for all sites across the calls. A second round of Delly was run using the ‘sites.bcf’ file to force genotyping across all sites. The resulting BCF files were merged using bcftools merge (-m id) and underwent downstream filtering (see below). Small variant calls made by GATK were merged and underwent joint genotyping using the GenomicsDBImport and GenotypeGVCFs functions.

### Linear Mapping and Variant Calling

All raw paired-end reads underwent a standard linear mapping and variant calling procedure to compare to calls using the pangenome approach. Raw reads underwent host contamination removal, trimming and QC as above. The resulting filtered FASTQ files were mapped to the 3D7 (v3) FASTA file using bwa (v0.7.17, -m flag enabled), followed by *de novo* variant calling using GATK (v.4.1.4.1), Delly (v1.5.0), Manta (v1.6.0) and Dysgu (v1.9.0) as above ^16,17,56^. Base Quality Score Recalibration was applied to BAM files using a high-quality *P. falciparum* genetic crosses dataset to correct systematic sequencing errors and instrument-specific base quality biases prior to variant calling ^7^; Variant Quality Score Recalibration was deliberately not applied to the benchmarking dataset to avoid introducing bias into the comparison, as it preferentially retains variants resembling the training data, which could unfairly favour or penalise calls from either the pangenome or linear reference-based approaches. Except for this, the pre-processing pipeline followed standard *P. falciparum* genomic processing practices ^53^. Variant calls were filtered using GATK SelectVariants and bcftools (v1.12). SNPs and indels were first separated using SelectVariants and filtered independently, applying the following thresholds: QUAL < 20 (low-confidence calls), MQ < 40 (poor mapping quality), QD < 2 (low quality-by-depth), FS > 60 (strand bias), BaseQRankSum or MQRankSum outside ±12.5 (rank sum outliers) and AF == 0 (invariant sites). Individual genotypes with fewer than five supporting reads (FMT/DP < 5) were set to missing, and sites with low total depth (INFO/DP < 5) were subsequently removed.

### Population Genetics Analysis

After processing samples using the linear and pangenome reference approaches, we investigated whether the pangenome method can be used for population genetics analysis. We first extracted biallelic SNPs (called using GATK from linear and surjected BAM files) from both datasets and used PLINK (v1.9) to produce a pairwise SNP distance matrix ^57^. We performed multidimensional scaling (MDS) (*K*= 5) of this matrix in R statistical software (v4.5.3) to compare population-level clustering from both methods. In addition, to assess whether population structure was driven by sequence shared with the linear reference, we recalculated the distance matrix and performed MDS restricted to variants representing sequence absent from 3D7. This included insertions whose alternate allele traversed graph nodes were not present in the 3D7 reference path, based on the allele traversal (AT) annotation output by VG Call ^18^. We next moved on to identifying any variants of interest at the population level. We first investigated any high-frequency variants across the entire dataset (MAF>0.05) in a list of drug-resistant candidate genes retrieved from the Malaria-Profiler database ^47^. We then performed Jost’s D analysis at two levels: for SVs (length ≥50 bp) and for small variants (length <50 bp). For small variants, we primarily focused on indels, as several global analyses have performed F_ST_ analysis of SNPs in the core genome at the global level previously ^25^. Jost’s D was calculated using a custom script with the normalised VCFs as input. Sites with >50% missing calls were removed from the analysis (https://github.com/cggh/scikit-allel). For each population, sites with the top 1% of Jost’s D values (99th percentile) were prioritised in the results.

For CNV detection, coverage was first estimated using mosdepth (v0.3.8) in 500 bp windows using the surjected BAM files as input ^58^. Even for linear mapping approaches, CNV calling is challenging, often relying on multiple lines of evidence including coverage-based analysis, detection of breakpoints, and manual inspection and curation. We therefore used a biologically informed approach for targeted CNV calling across *pfmdr1*, employing precise amplification breakpoint coordinates from the Pf8 dataset (https://pf8-release.cog.sanger.ac.uk/Pf8_tandem_duplication_breakpoints.tsv) ^4^. For each of the *pfmdr1* breakpoint patterns, flanking regions of approximately 20 kb upstream and downstream of the target amplification region were defined. Weighted average coverage was extracted from mosdepth output for three regions per pattern: the target breakpoint region, the left flanking region and the right flanking region. Per-sample normalisation was performed by calculating a flanking baseline as the mean of left and right flanking coverage, with the coverage ratio defined as target coverage divided by this baseline, controlling for sample-specific sequencing depth variation. Two quality filters were applied: patterns where mean flank coverage fell below 5x were excluded (789 samples in total remained), and patterns with greater than 1.5-fold left-to-right flank imbalance were flagged and excluded as potentially artefactual.

CNVs were classified as high confidence (ratio ≥2.0), medium confidence (ratio ≥1.5), or low confidence (ratio ≥1.3) amplifications, with calls aggregated to gene level requiring at least one medium confidence pattern call. Concordance of high-confidence pangenome-derived *pfmdr1* calls was assessed against expert-curated Pf8 amplification calls ^4^. To characterise smaller-scale CNVs, including at the structurally complex *pfgch1* locus, indel sequences were extracted using a custom script with alignments generated by MAFFT (v7.536) and BLASTed against the 3D7 genome using BLAST (v2.16.0) to identify gene-level duplications ^59,60^. CNVs at *pfgch1* were confirmed using normalised coverage plots of the linear-mapped BAM files for the 12 reference assembly sequences aligned to the 3D7 reference.

## Data availability

This publication uses MalariaGEN data as described in ‘Pf8: an open dataset of *Plasmodium falciparum* genome variation in 33,325 worldwide samples.’ MalariaGEN et al, Wellcome Open Research 2025, **10**:325 https://doi.org/10.12688/wellcomeopenres.24031.1. A full list of accessions is available in Supplementary Table 11. No new samples were sequenced as a part of this project. PfPan is available in a dedicated repository: https://genomics.lshtm.ac.uk/data/ and via Zenodo: 10.5281/zenodo.23218483.

## Code availability

The code used is available from open-access packages mentioned in **Methods**. For a detailed overview, please see the dedicated GitHub repository: https://github.com/NinaMercedes/PfPan.

## Acknowledgements

N/A

## Contributions

NB conceptualised and performed all analyses and drafted the manuscript. TGC and SC contributed to conceptualisation, provided supervision, and reviewed the results and the manuscript. SC and JT assisted with dataset curation. All authors reviewed and approved the final manuscript.

## References

1 Gardner, M. J. et al. Genome sequence of the human malaria parasite Plasmodium falciparum. Nature 419, 498–511 (2002). 10.1038/nature01097

2 Walliker, D. et al. Genetic analysis of the human malaria parasite Plasmodium falciparum. Science 236, 1661–1666 (1987). 10.1126/science.3299700

3 Bohme, U., Otto, T. D., Sanders, M., Newbold, C. I. & Berriman, M. Progression of the canonical reference malaria parasite genome from 2002-2019. Wellcome Open Res 4, 58 (2019). 10.12688/wellcomeopenres.15194.2

4 MalariaGEN, P. f. C. P., et al. Pf8: an open dataset of Plasmodium falciparum genome variation in 33,325 worldwide samples [version 1; peer review: 2 approved]. Wellcome Open Research 10 (2025). 10.12688/wellcomeopenres.24031.1

5 Bruske, E., Otto, T. D. & Frank, M. Whole genome sequencing and microsatellite analysis of the Plasmodium falciparum E5 NF54 strain show that the var, rifin and stevor gene families follow Mendelian inheritance. Malar J 17, 376 (2018). 10.1186/s12936-018-2503-2

6 Otto, T. D. et al. Long read assemblies of geographically dispersed Plasmodium falciparum isolates reveal highly structured subtelomeres. Wellcome Open Res 3, 52 (2018). 10.12688/wellcomeopenres.14571.1

7 Miles, A. et al. Indels, structural variation, and recombination drive genomic diversity in Plasmodium falciparum. Genome Res 26, 1288–1299 (2016). 10.1101/gr.203711.115

8 Liao, W.-W. et al. A draft human pangenome reference. Nature 617, 312–324 (2023). 10.1038/s41586-023-05896-x

9 Loegler, V. et al. From genotype to phenotype with 1,086 near telomere-to-telomere yeast genomes. Nature 648, 649–658 (2025). 10.1038/s41586-025-09637-0

10 Jiao, C. et al. Pan-genome bridges wheat structural variations with habitat and breeding. Nature 637, 384–393 (2025). 10.1038/s41586-024-08277-0

11 Wang, K. et al. The Chicken Pan-Genome Reveals Gene Content Variation and a Promoter Region Deletion in IGF2BP1 Affecting Body Size. Molecular Biology and Evolution 38, 5066–5081 (2021). 10.1093/molbev/msab231

12 Rice, E. S. et al. A pangenome graph reference of 30 chicken genomes allows genotyping of large and complex structural variants. BMC Biol 21, 267 (2023). 10.1186/s12915-023-01758-0

13 Arshad, F. et al. A comprehensive water buffalo pangenome reveals extensive structural variation linked to population-specific signatures of selection. GigaScience 14, giaf099 (2025). 10.1093/gigascience/giaf099

14 Letcher, B., Hunt, M. & Iqbal, Z. Gramtools enables multiscale variation analysis with genome graphs. Genome Biol 22, 259 (2021). 10.1186/s13059-021-02474-0

15 Ravenhall, M. et al. An analysis of large structural variation in global Plasmodium falciparum isolates identifies a novel duplication of the chloroquine resistance associated gene. Scientific Reports 9, 8287 (2019). 10.1038/s41598-019-44599-0

16 McKenna, A. et al. The Genome Analysis Toolkit: a MapReduce framework for analyzing next-generation DNA sequencing data. Genome Res 20, 1297–1303 (2010). 10.1101/gr.107524.110

17 Rausch, T. et al. DELLY: structural variant discovery by integrated paired-end and split-read analysis. Bioinformatics 28, i333–i339 (2012). 10.1093/bioinformatics/bts378

18 Hickey, G. et al. Genotyping structural variants in pangenome graphs using the vg toolkit. Genome Biol 21, 35 (2020). 10.1186/s13059-020-1941-7

19 Ghumra, A. et al. Induction of strain-transcending antibodies against Group A PfEMP1 surface antigens from virulent malaria parasites. PLoS Pathog 8, e1002665 (2012). 10.1371/journal.ppat.1002665

20 Mu, J. et al. Recombination hotspots and population structure in Plasmodium falciparum. PLoS Biol 3, e335 (2005). 10.1371/journal.pbio.0030335

21 Fola, A. A. et al. Plasmodium falciparum resistant to artemisinin and diagnostics have emerged in Ethiopia. Nature Microbiology 8, 1911–1919 (2023). 10.1038/s41564-023-01461-4

22 Gamboa, D. et al. A large proportion of P. falciparum isolates in the Amazon region of Peru lack pfhrp2 and pfhrp3: implications for malaria rapid diagnostic tests. PLoS One 5, e8091 (2010). 10.1371/journal.pone.0008091

23 Góes, L. et al. Evaluation of Histidine-Rich Proteins 2 and 3 Gene Deletions in Plasmodium falciparum in Endemic Areas of the Brazilian Amazon. International Journal of Environmental Research and Public Health 18 (2021).

24 Kamaliddin, C. et al. A Countrywide Survey of hrp2/3 Deletions and kelch13 Mutations Co-occurrence in Ethiopia. J Infect Dis 230, e1394–e1401 (2024). 10.1093/infdis/jiae373

25 Billows, N. et al. Global-scale population genetic analysis of Plasmodium falciparum identifies region-specific patterns of malaria parasite adaptation. Nature Communications (2026). 10.1038/s41467-026-73006-2

26 Eksi, S. et al. Plasmodium falciparum gametocyte development 1 (Pfgdv1) and gametocytogenesis early gene identification and commitment to sexual development. PLoS Pathog 8, e1002964 (2012). 10.1371/journal.ppat.1002964

27 Sallicandro, P. et al. Repetitive sequences upstream of the pfg27/25 gene determine polymorphism in laboratory and natural lines of Plasmodium falciparum. Molecular and Biochemical Parasitology 110, 247–257 (2000). 10.1016/S0166-6851(00)00274-7

28 Scherf, A. et al. Gene inactivation of Pf11-1 of Plasmodium falciparum by chromosome breakage and healing: identification of a gametocyte-specific protein with a potential role in gametogenesis. Embo j 11, 2293–2301 (1992). 10.1002/j.1460-2075.1992.tb05288.x

29 Lim, M. Y.-X. et al. UDP-galactose and acetyl-CoA transporters as Plasmodium multidrug resistance genes. Nature Microbiology 1, 16166 (2016). 10.1038/nmicrobiol.2016.166

30 Lee, C. et al. Pf-HaploAtlas: an interactive web app for spatiotemporal analysis of Plasmodium falciparum genes. Bioinformatics 40, btae673 (2024). 10.1093/bioinformatics/btae673

31 McHugh, E. et al. A repeat sequence domain of the ring-exported protein-1 of Plasmodium falciparum controls export machinery architecture and virulence protein trafficking. Mol Microbiol 98, 1101–1114 (2015). 10.1111/mmi.13201

32 Josling, G. A. et al. Dissecting the role of PfAP2-G in malaria gametocytogenesis. Nature Communications 11, 1503 (2020). 10.1038/s41467-020-15026-0

33 Singer, M., Kanatani, S., Castillo, S. G., Frischknecht, F. & Sinnis, P. The Plasmodium circumsporozoite protein. Trends Parasitol 40, 1124–1134 (2024). 10.1016/j.pt.2024.10.017

34 Amato, R. et al. Origins of the current outbreak of multidrug-resistant malaria in southeast Asia: a retrospective genetic study. Lancet Infect Dis 18, 337–345 (2018). 10.1016/s1473-3099(18)30068-9

35 Nair, S. et al. Adaptive Copy Number Evolution in Malaria Parasites. PLOS Genetics 4, e1000243 (2008). 10.1371/journal.pgen.1000243

36 Ma, W. & Chaisson, M. J. P. Genotyping sequence-resolved copy number variation using pangenomes reveals paralog-specific global diversity and expression divergence of duplicated genes. Nature Genetics 57, 2909–2919 (2025). 10.1038/s41588-025-02346-4

37 Liu, S. et al. Direct long-read visualization reveals hidden variation in GCH1 gene copy number and precise expansion steps. BMC Genomics 26, 671 (2025). 10.1186/s12864-025-11859-5

38 Zhu, S. J., Almagro-Garcia, J. & McVean, G. Deconvolution of multiple infections in Plasmodium falciparum from high throughput sequencing data. Bioinformatics 34, 9–15 (2018). 10.1093/bioinformatics/btx530

39 Poplin, R. et al. A universal SNP and small-indel variant caller using deep neural networks. Nature Biotechnology 36, 983–987 (2018). 10.1038/nbt.4235

40 Hickey, G. et al. Pangenome graph construction from genome alignments with Minigraph-Cactus. Nat Biotechnol 42, 663–673 (2024). 10.1038/s41587-023-01793-w

41 Parmigiani, L., Garrison, E., Stoye, J., Marschall, T. & Doerr, D. Panacus: fast and exact pangenome growth and core size estimation. Bioinformatics 40, btae720 (2024). 10.1093/bioinformatics/btae720

42 Guarracino, A., Heumos, S., Nahnsen, S., Prins, P. & Garrison, E. ODGI: understanding pangenome graphs. Bioinformatics 38, 3319–3326 (2022). 10.1093/bioinformatics/btac308

43 Li, H. Minimap2: pairwise alignment for nucleotide sequences. Bioinformatics 34, 3094–3100 (2018). 10.1093/bioinformatics/bty191

44 Heller, D. & Vingron, M. SVIM-asm: structural variant detection from haploid and diploid genome assemblies. Bioinformatics 36, 5519–5521 (2021). 10.1093/bioinformatics/btaa1034

45 Danecek, P. et al. Twelve years of SAMtools and BCFtools. GigaScience 10, giab008 (2021). 10.1093/gigascience/giab008

46 English, A. C., Menon, V. K., Gibbs, R. A., Metcalf, G. A. & Sedlazeck, F. J. Truvari: refined structural variant comparison preserves allelic diversity. Genome Biology 23, 271 (2022). 10.1186/s13059-022-02840-6

47 Phelan, J. E. et al. Rapid profiling of Plasmodium parasites from genome sequences to assist malaria control. Genome Med 15, 96 (2023). 10.1186/s13073-023-01247-7

48 World Health Organization. Malaria: Artemisinin partial resistance. https://www.who.int/news-room/questions-and-answers/item/artemisinin-resistance. (2025).

49 World Health, O. Compendium of molecular markers for antimalarial drug resistance. (2025).

50 Beyer, W. et al. Sequence tube maps: making graph genomes intuitive to commuters. Bioinformatics 35, 5318–5320 (2019). 10.1093/bioinformatics/btz597

51 Wick, R. R., Schultz, M. B., Zobel, J. & Holt, K. E. Bandage: interactive visualization of de novo genome assemblies. Bioinformatics 31, 3350–3352 (2015). 10.1093/bioinformatics/btv383

52 Auburn, S. et al. Characterization of Within-Host Plasmodium falciparum Diversity Using Next-Generation Sequence Data. PLOS ONE 7, e32891 (2012). 10.1371/journal.pone.0032891

53 Abdel Hamid, M. M., et al. Pf7: an open dataset of Plasmodium falciparum genome variation in 20,000 worldwide samples. Wellcome Open Res 8, 22 (2023). 10.12688/wellcomeopenres.18681.1

54 Constantinides, B., Hunt, M. & Crook, D. W. Hostile: accurate decontamination of microbial host sequences. Bioinformatics 39, btad728 (2023). 10.1093/bioinformatics/btad728

55 Chen, S., Zhou, Y., Chen, Y. & Gu, J. fastp: an ultra-fast all-in-one FASTQ preprocessor. Bioinformatics 34, i884–i890 (2018). 10.1093/bioinformatics/bty560

56 Li, H. & Durbin, R. Fast and accurate short read alignment with Burrows-Wheeler transform. Bioinformatics 25, 1754–1760 (2009). 10.1093/bioinformatics/btp324

57 Purcell, S. et al. PLINK: a tool set for whole-genome association and population-based linkage analyses. Am J Hum Genet 81, 559–575 (2007). 10.1086/519795

58 Pedersen, B. S. & Quinlan, A. R. Mosdepth: quick coverage calculation for genomes and exomes. Bioinformatics 34, 867–868 (2018). 10.1093/bioinformatics/btx699

59 Katoh, K. & Standley, D. M. MAFFT multiple sequence alignment software version 7: improvements in performance and usability. Mol Biol Evol 30, 772–780 (2013). 10.1093/molbev/mst010

60 Altschul, S. F., Gish, W., Miller, W., Myers, E. W. & Lipman, D. J. Basic local alignment search tool. Journal of Molecular Biology 215, 403–410 (1990). 10.1016/S0022-2836(05)80360-2

