## Supplementary Figure for "A *Plasmodium falciparum* Pangenome Resource to drive Structural Variant Discovery and assist Malaria Control"

* Joint authors

Prof. Susana Campino, LSHTM

Prof. Taane G. Clark, LSHTM

**Supplementary Figures**

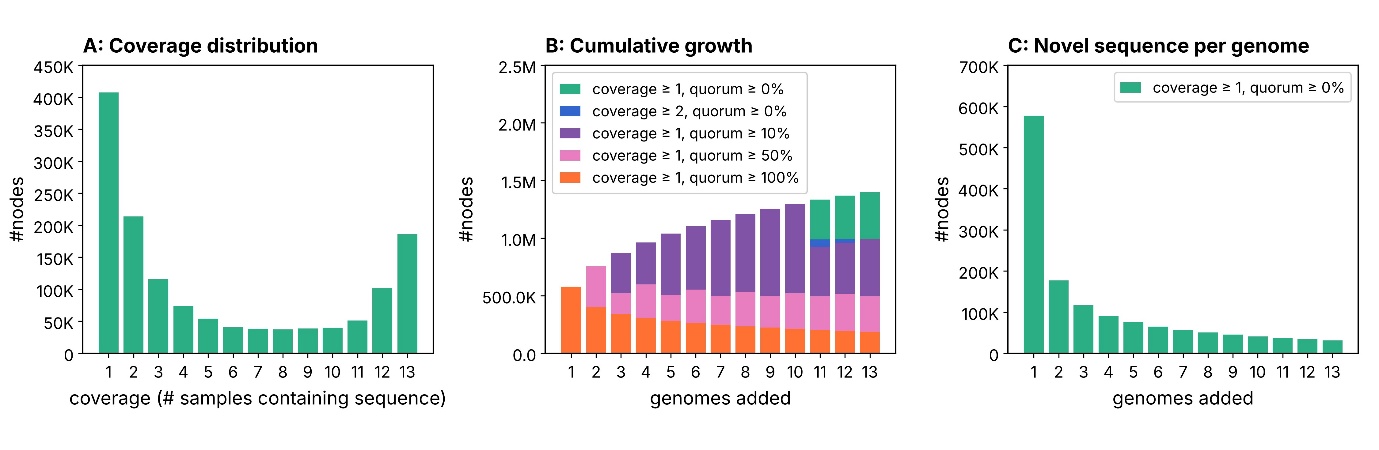

**Supplementary Figure 1. Pangenome growth statistics (nodes).** **(A)** Distribution of node coverage: the number of nodes present in exactly *n* of the 13 genomes (n = 1, private; n = 13, core). **(B)** Cumulative number of pangenome nodes as genomes are added one at a time, shown across five coverage/quorum thresholds. **(C)** Novel nodes contributed by each additional genome.

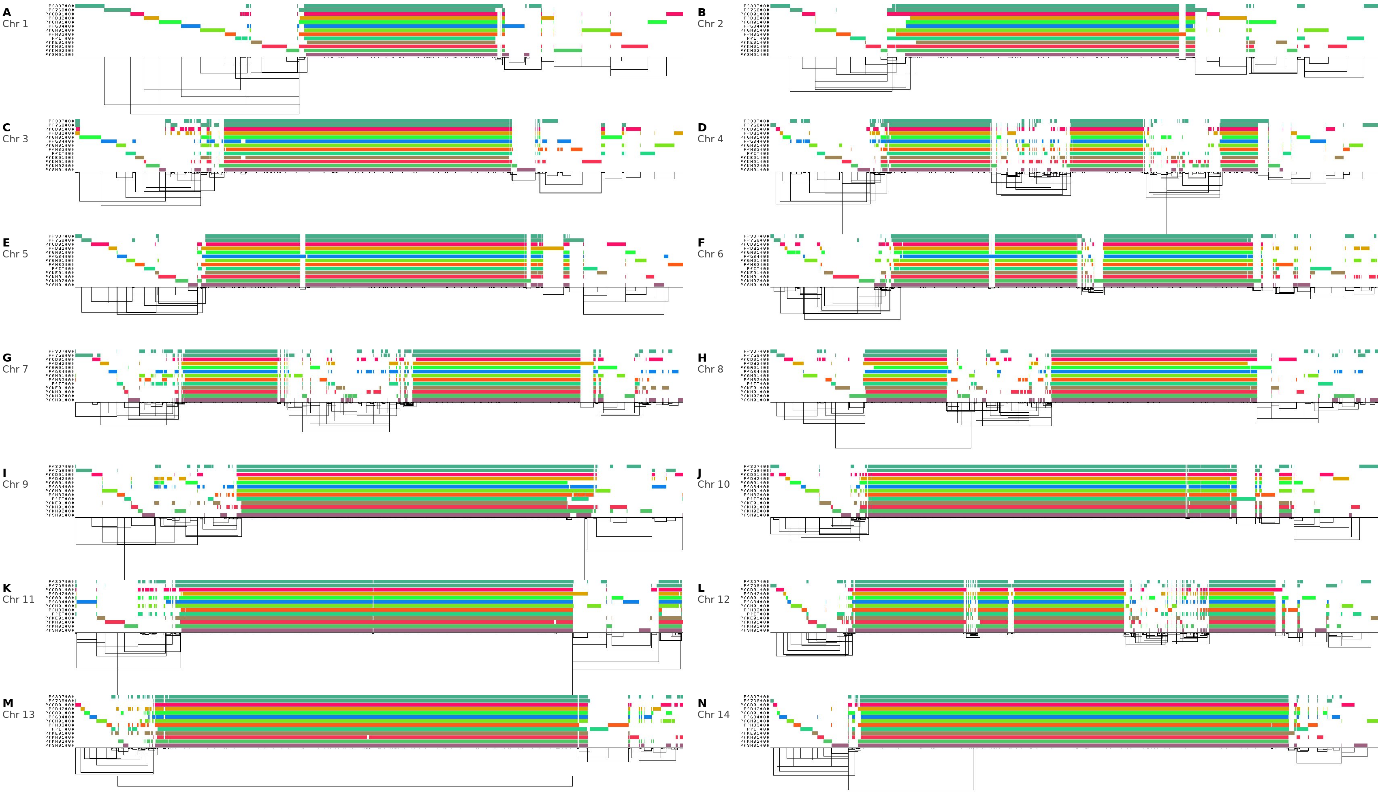

**Supplementary Figure 2. ODGI visualisation of pangenome alignments across all 14 chromosomes.** ODGI visualisation of pangenome alignments across all 14 *P. falciparum* chromosomes (panels A–N), illustrating core genome continuity, internal hypervariable regions, and subtelomeric boundaries across the 13 reference strains.

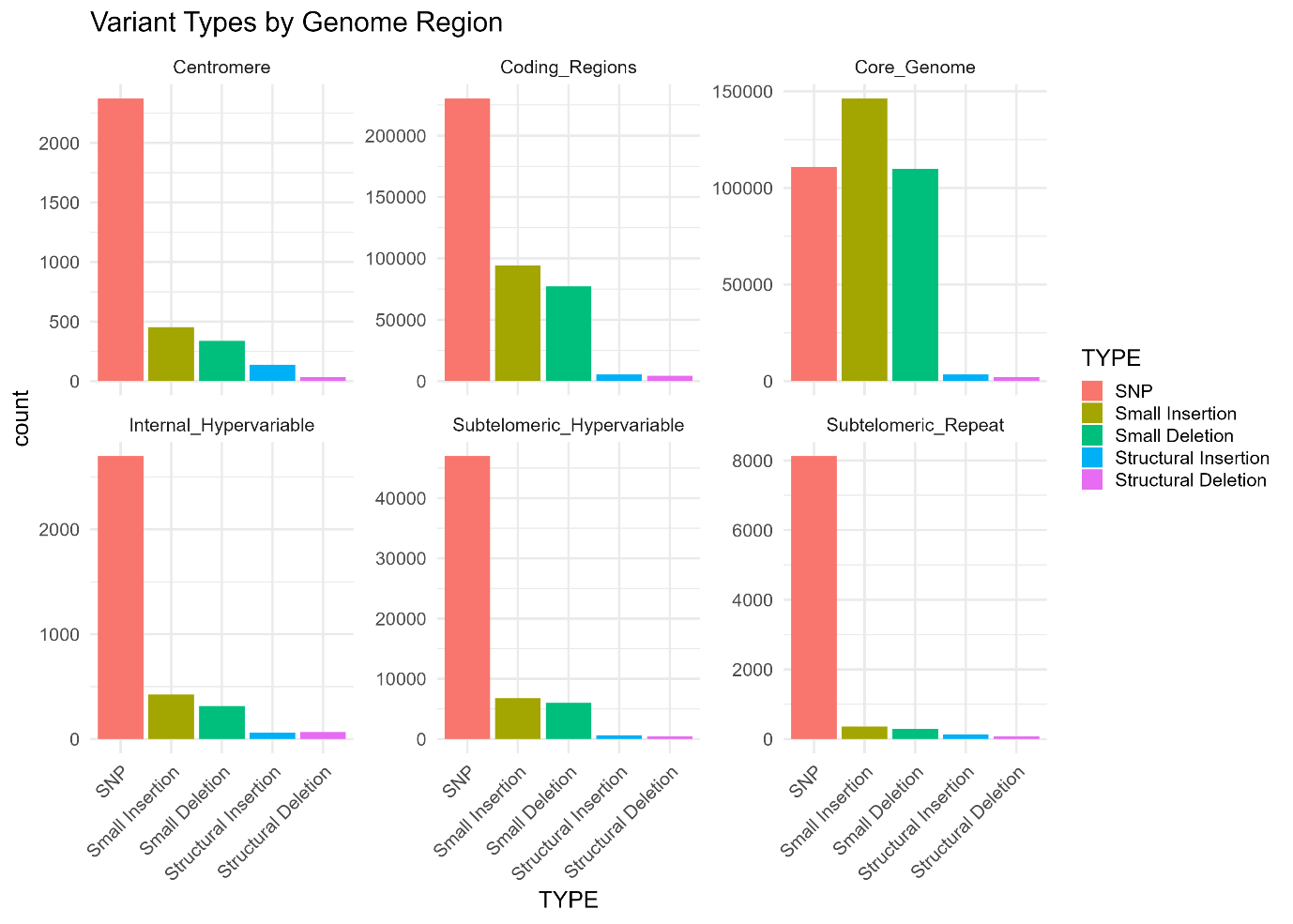

**Supplementary Figure 3. Variant type distributions by genomic region.** Variant type distributions stratified by genomic region (core, coding, centromeric, internal hypervariable, and subtelomeric hypervariable) across SNPs, insertions, deletions, structural insertions, and structural deletions.

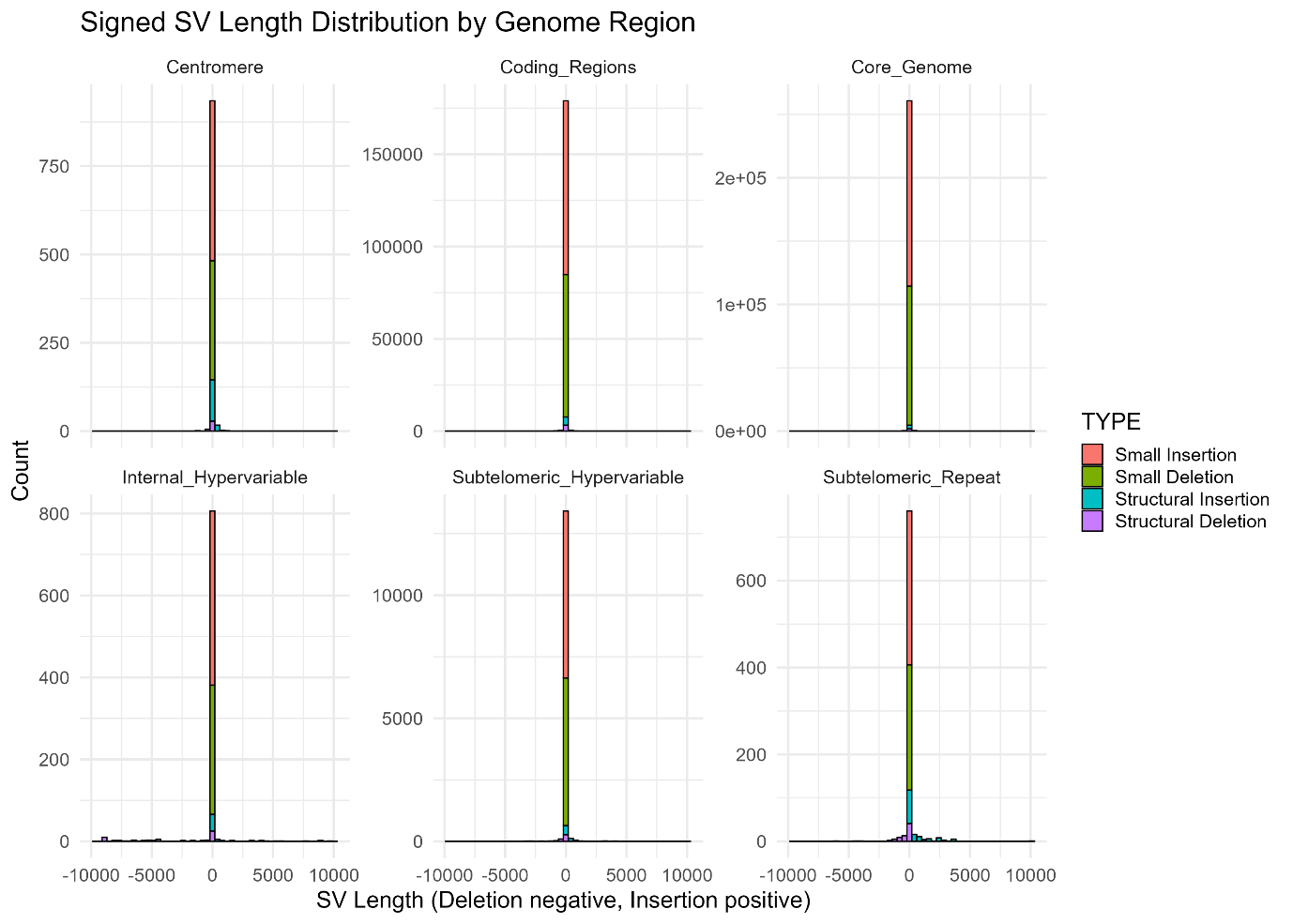

**Supplementary Figure 4. Variant length distributions by class.** Length distributions of variant classes across SNPs, insertions, deletions, structural insertions, and structural deletions identified across the PfPan13 reference genomes.

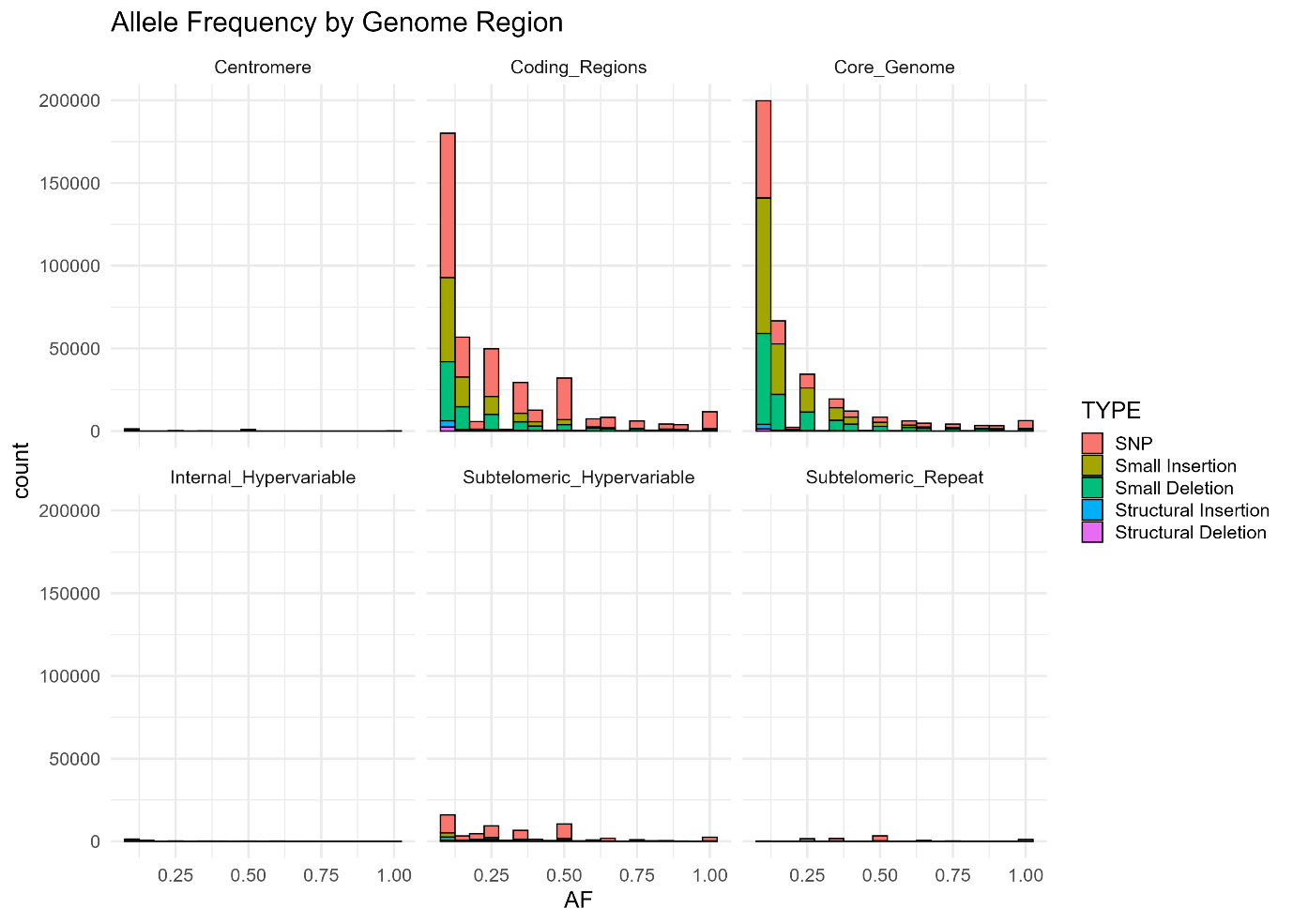

**Supplementary Figure 5. Allele frequency distributions by genomic region and variant type.** Allele frequency distributions stratified by genomic region (core, coding, centromeric, internal hypervariable, and subtelomeric hypervariable) and variant type across PfPan.

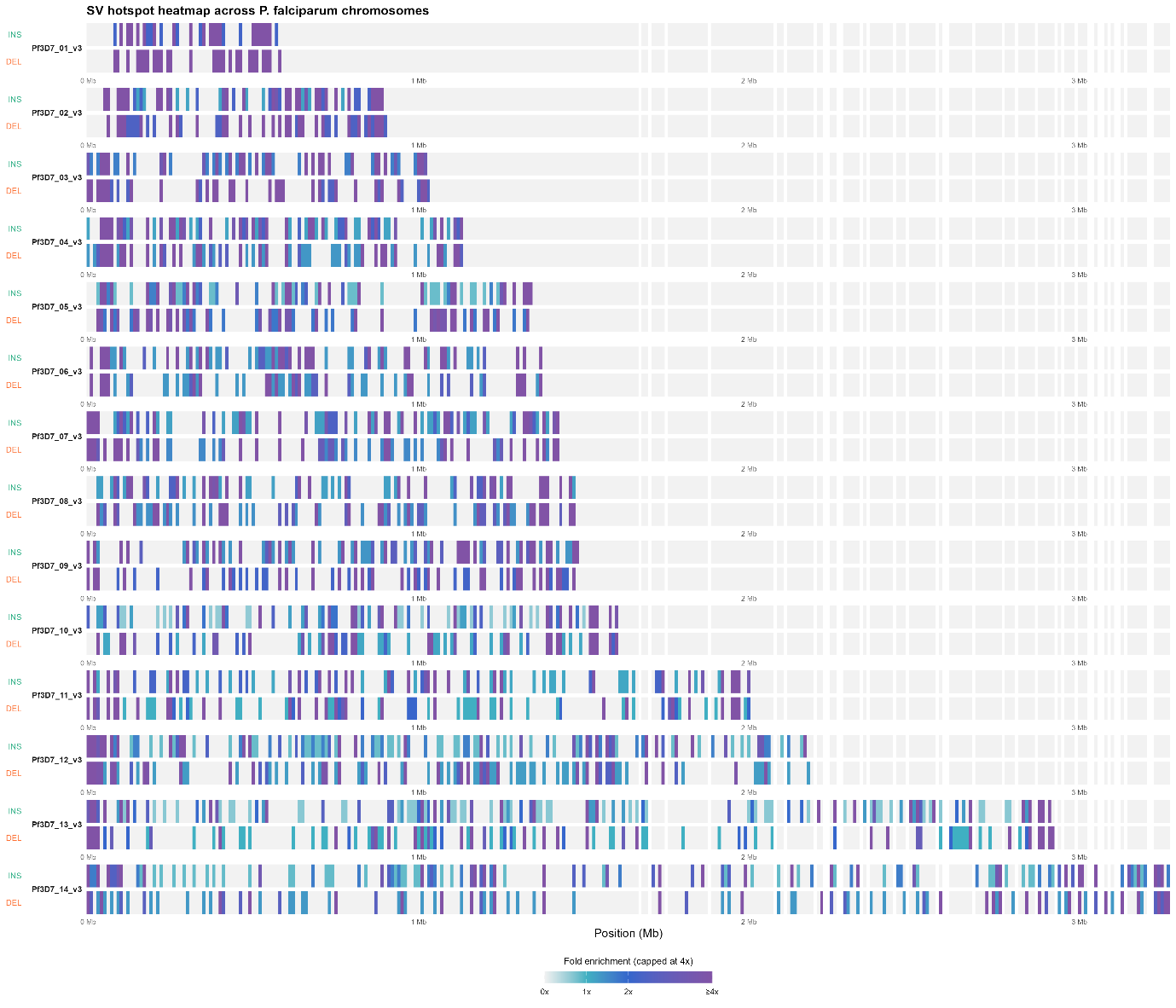

**Supplementary Figure 6. Genome-wide structural variant enrichment heatmaps.** Genome-wide enrichment of structural variants (≥50 bp) in 10 kb windows across all 14 chromosomes relative to Pf3D7 coordinates, shown as heatmaps for insertions (top) and deletions (bottom).

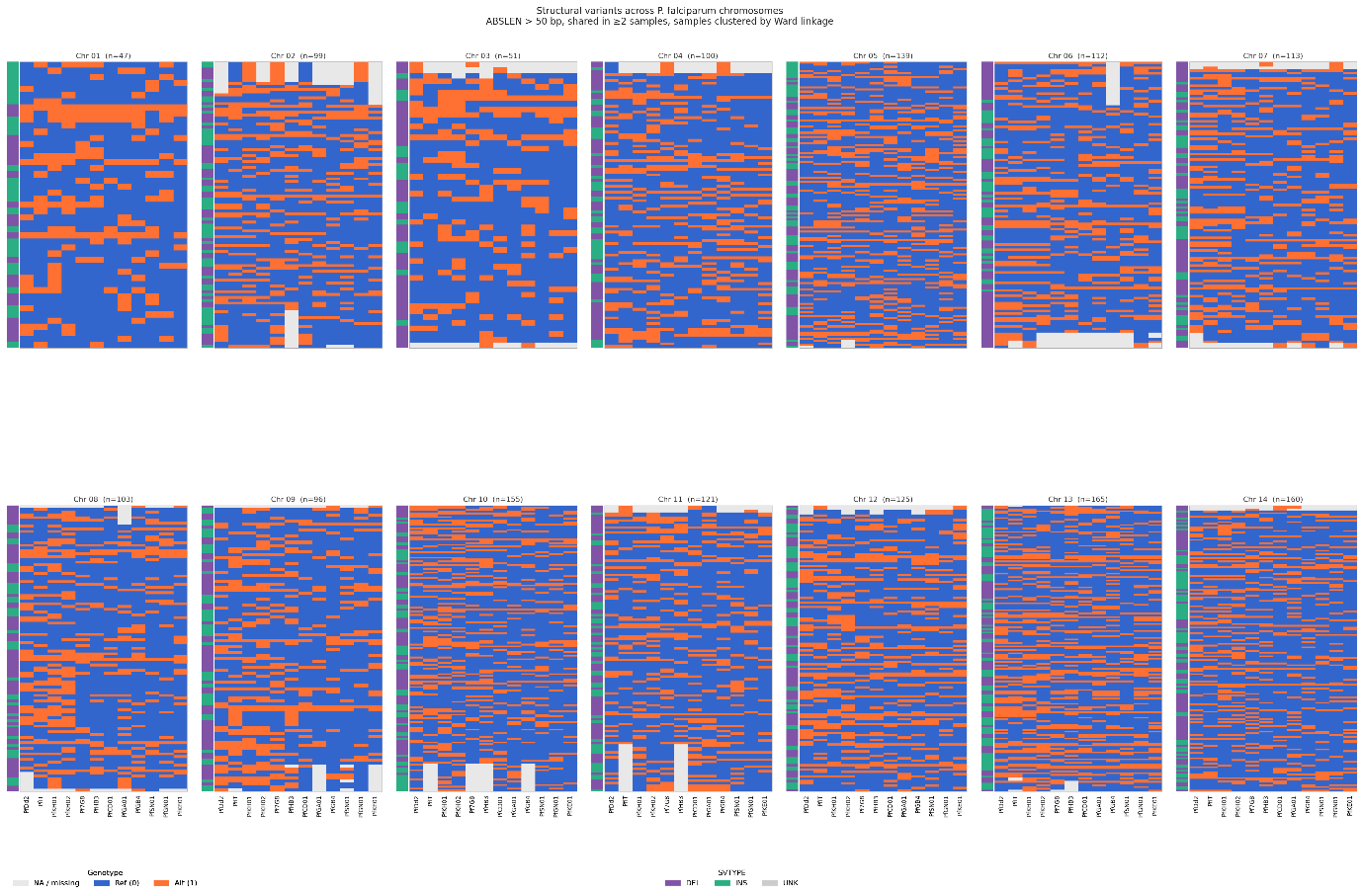

**Supplementary Figure 7. Per-chromosome structural variant haplotype heatmaps.** Per-chromosome heatmaps of structural variant haplotypes (structural insertions and deletions) across the 13 PfPan reference strains (x-axis), with cells coloured by alternate versus reference allele state and strains clustered by Ward linkage. Genomic region annotations are indicated by the coloured panel on the left.

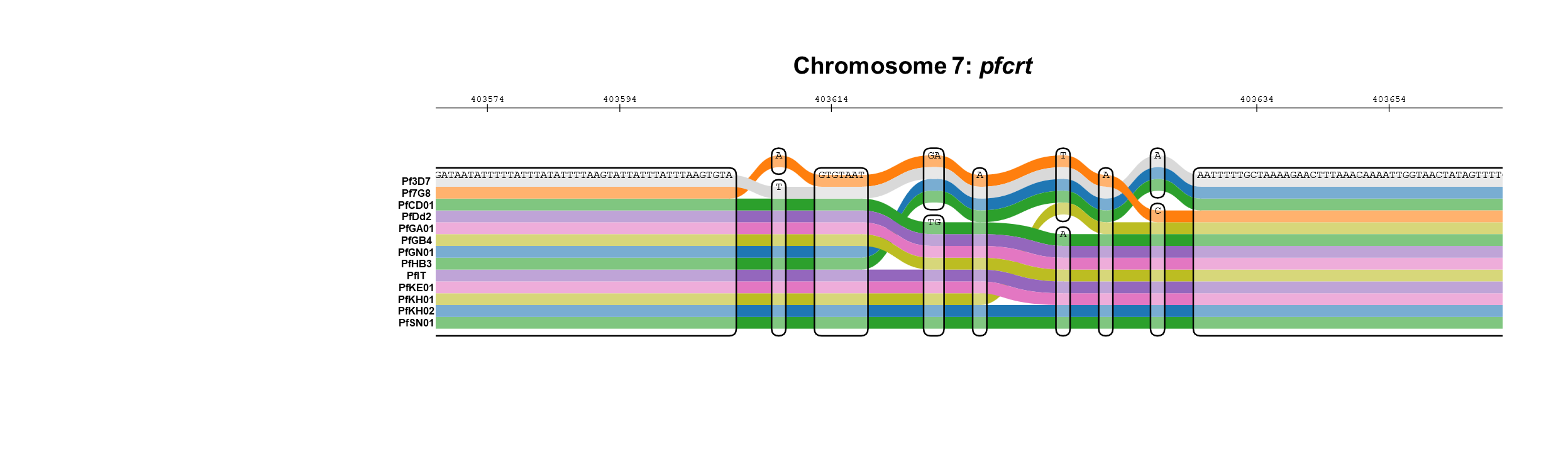

**Supplementary Figure 8. Sequence tube map of *pfcrt*.** Sequence tube map of the *pfcrt* illustrating haplotype diversity across PfPan reference strains, including the CVIET haplotype associated with chloroquine resistance.

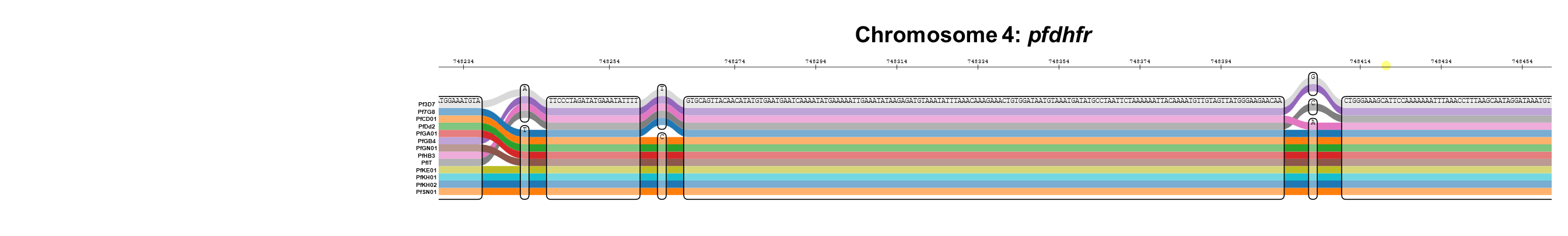

**Supplementary Figure 9. Sequence tube map of *pfdhfr*.** Sequence tube map of *pfdhfr* illustrating haplotype diversity across PfPan reference strains, including the triple mutant haplotype associated with pyrimethamine resistance.

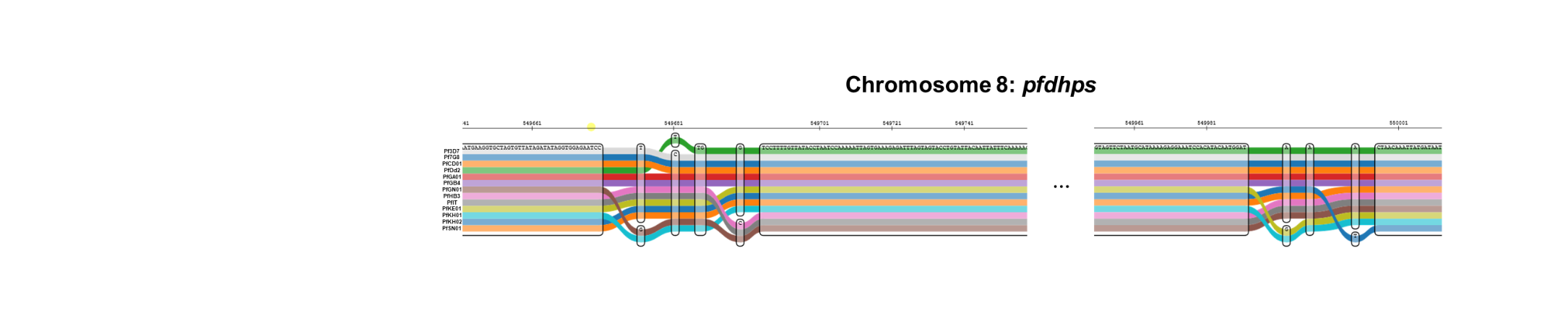

**Supplementary Figure 10. Sequence tube map of the *pfdhps* locus.** Sequence tube map of the *pfdhps* locus illustrating haplotype diversity across PfPan reference strains.

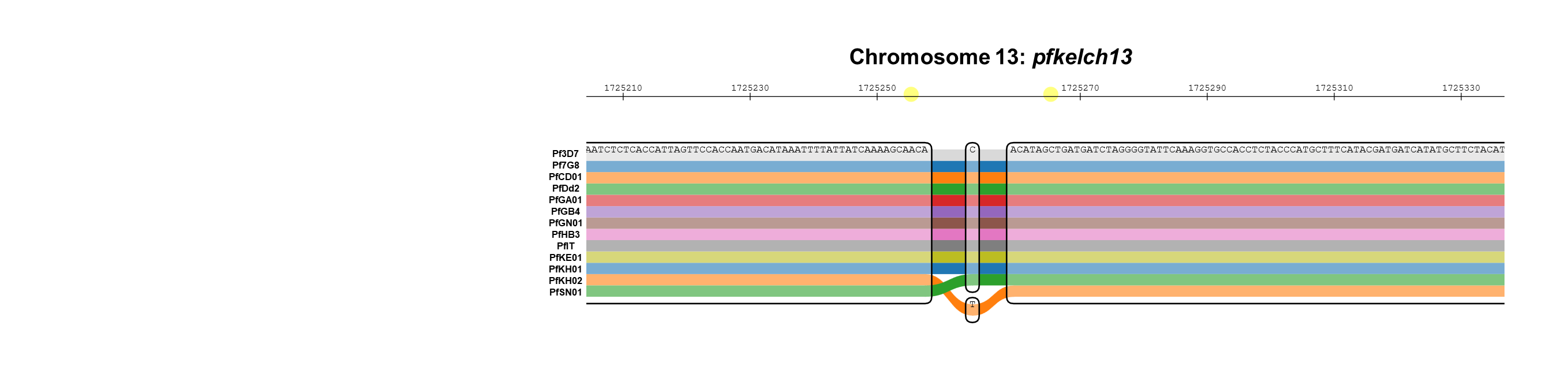
**Supplementary Figure 11. Sequence tube map of *pfkelch13*.** Sequence tube map of *pfkelch13*, highlighting the C580Y artemisinin partial resistance mutation present exclusively in the Cambodian reference strain KH02.

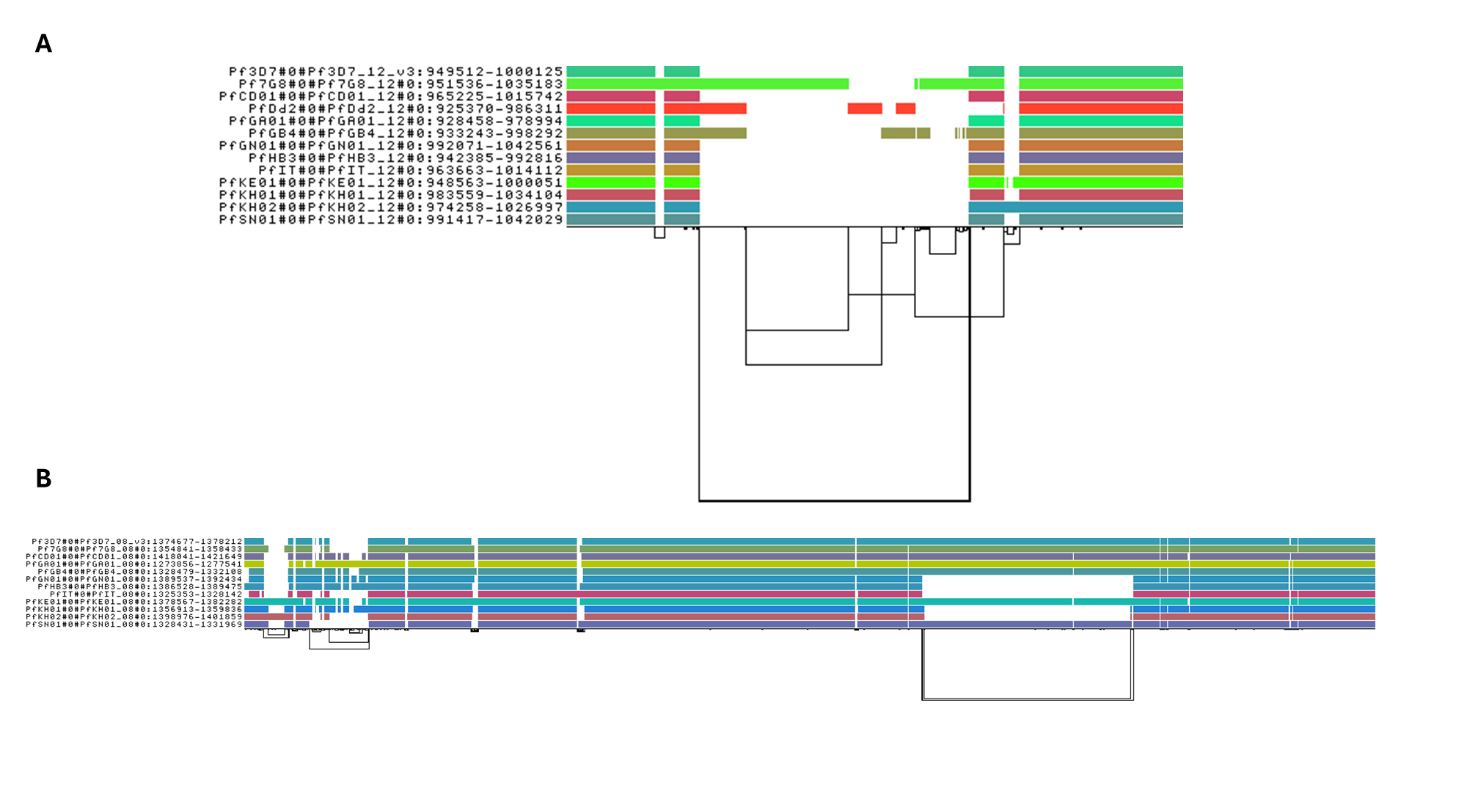

**Supplementary Figure 12. ODGI visualisations of the *pfgch1* and *pfhrp2* loci.** ODGI visualisations of *pfgch1* (A) and *pfhrp2* (B), illustrating structural complexity and sequence divergence across PfPan reference strains.

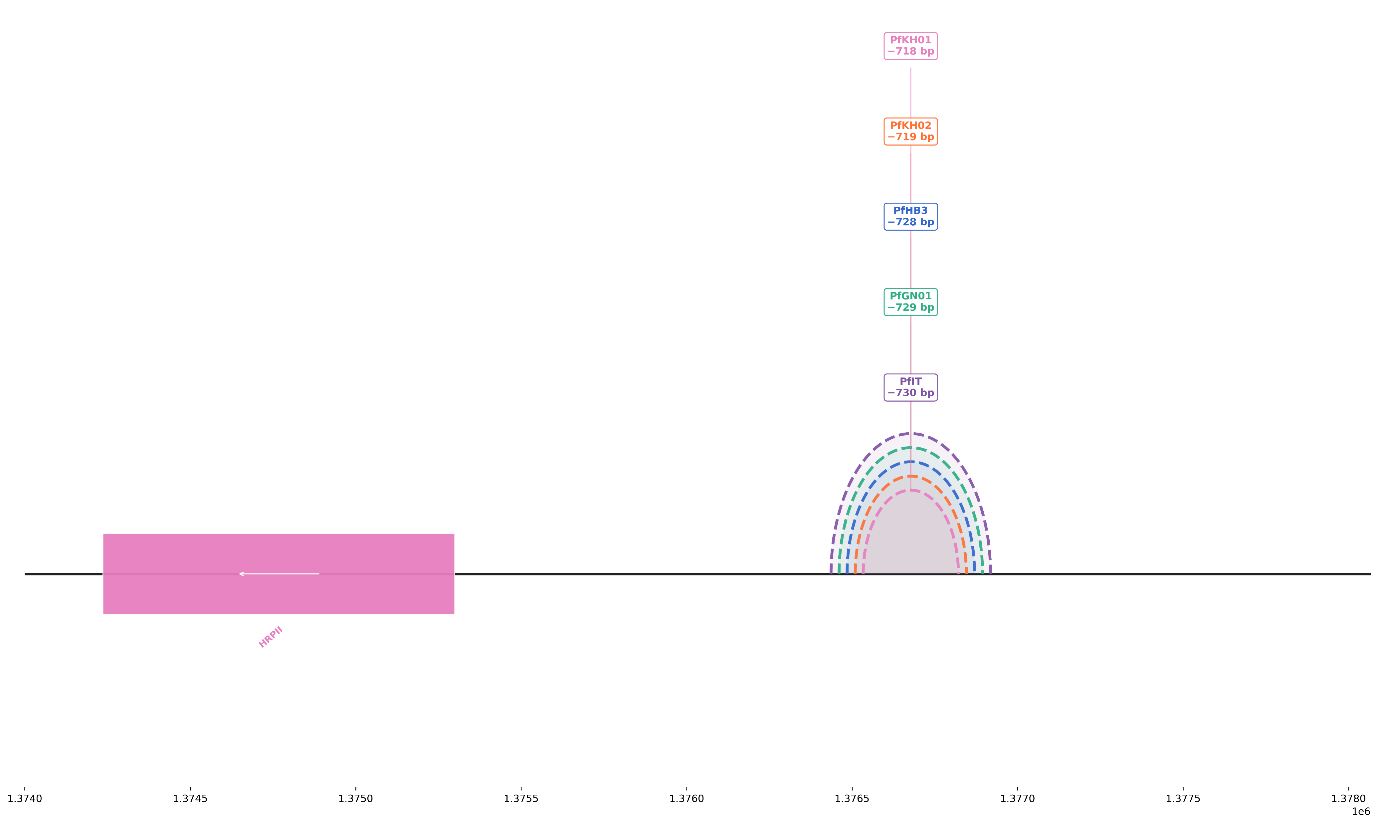

**Supplementary Figure 13. Large-scale *pfhrp2* upstream deletions across reference strains.** Arc diagrams of *pfhrp2* showing large-scale upstream promoter deletions of variable size across reference strains KH02, IT, GN01, HB3, and KH01.

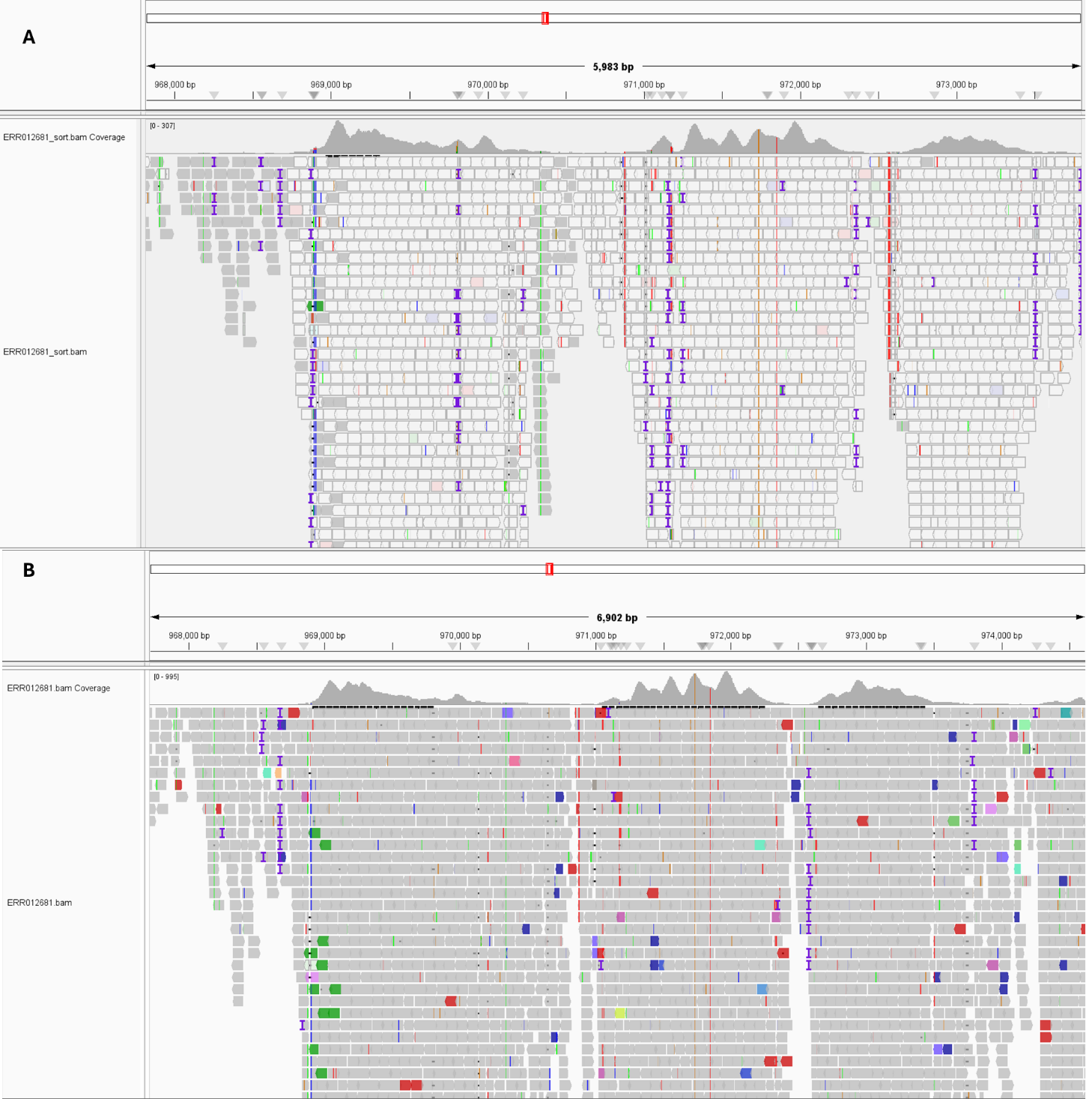

**Supplementary Figure 14. Comparative IGV visualisation of *pfgch1* copy number variant mapping.** IGV screenshots of the *pfgch1*region for isolate ERR012681, a sample with known GCH1 and MDR1 copy number variants, comparing pangenome-surjected alignments (A) and linear reference-mapped alignments (B). Lighter shading in the pangenome-surjected BAM reflects reduced mapping quality arising from coordinate surjection at this structurally complex locus, whereas linear mappings show elevated and high-quality coverage consistent with copy number amplification.

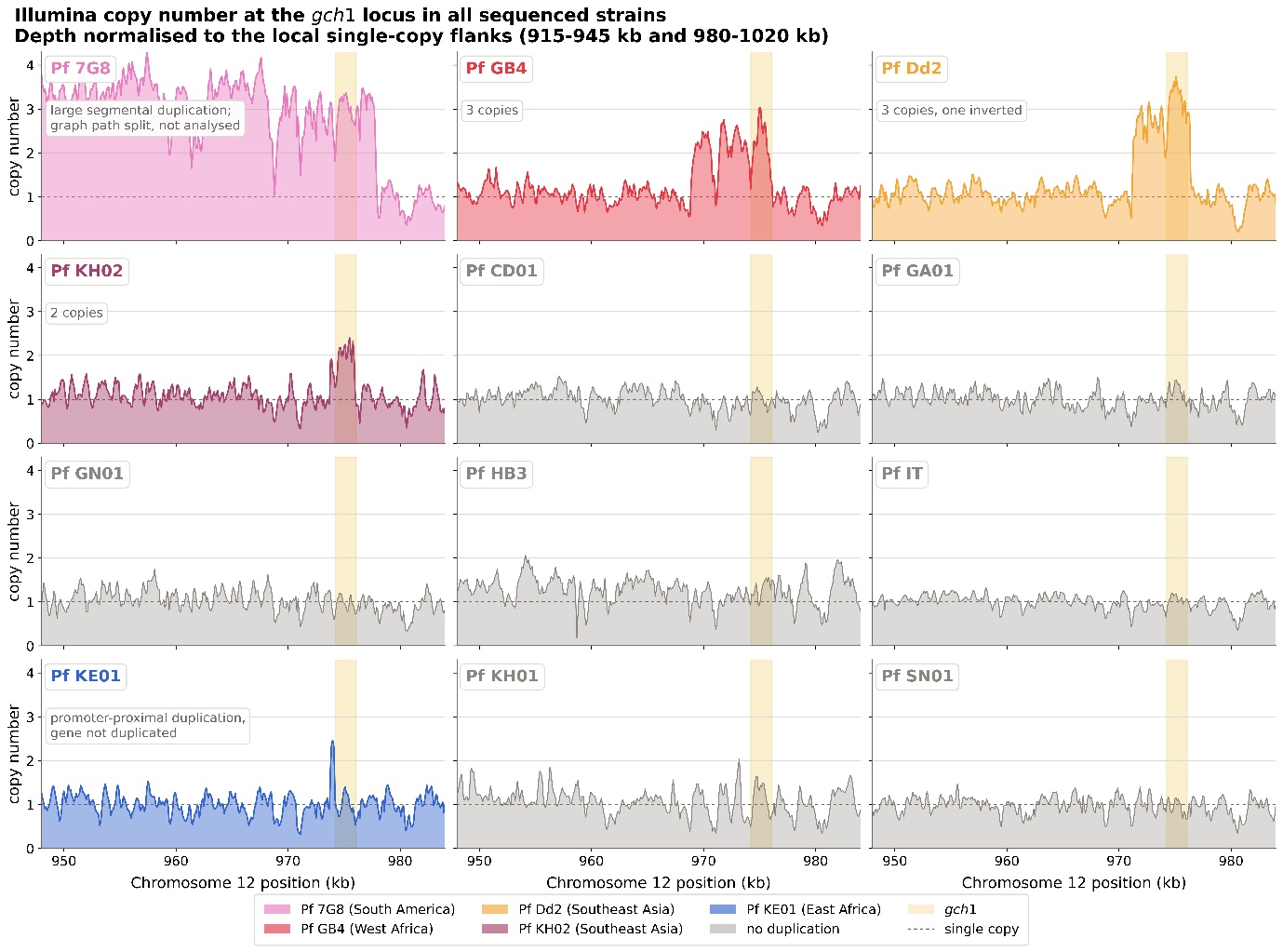
 **Supplementary Figure 15. Copy number variation across *pfgch1.*** Filled areas show Illumina read depth in 100 bp bins, normalised to the median depth of the local single-copy flanks (915,000–945,000 and 980,000–1,020,000 bp). The dashed line marks single copy number and the shaded band highlights *pfgch1* (974,225–976,097 bp).

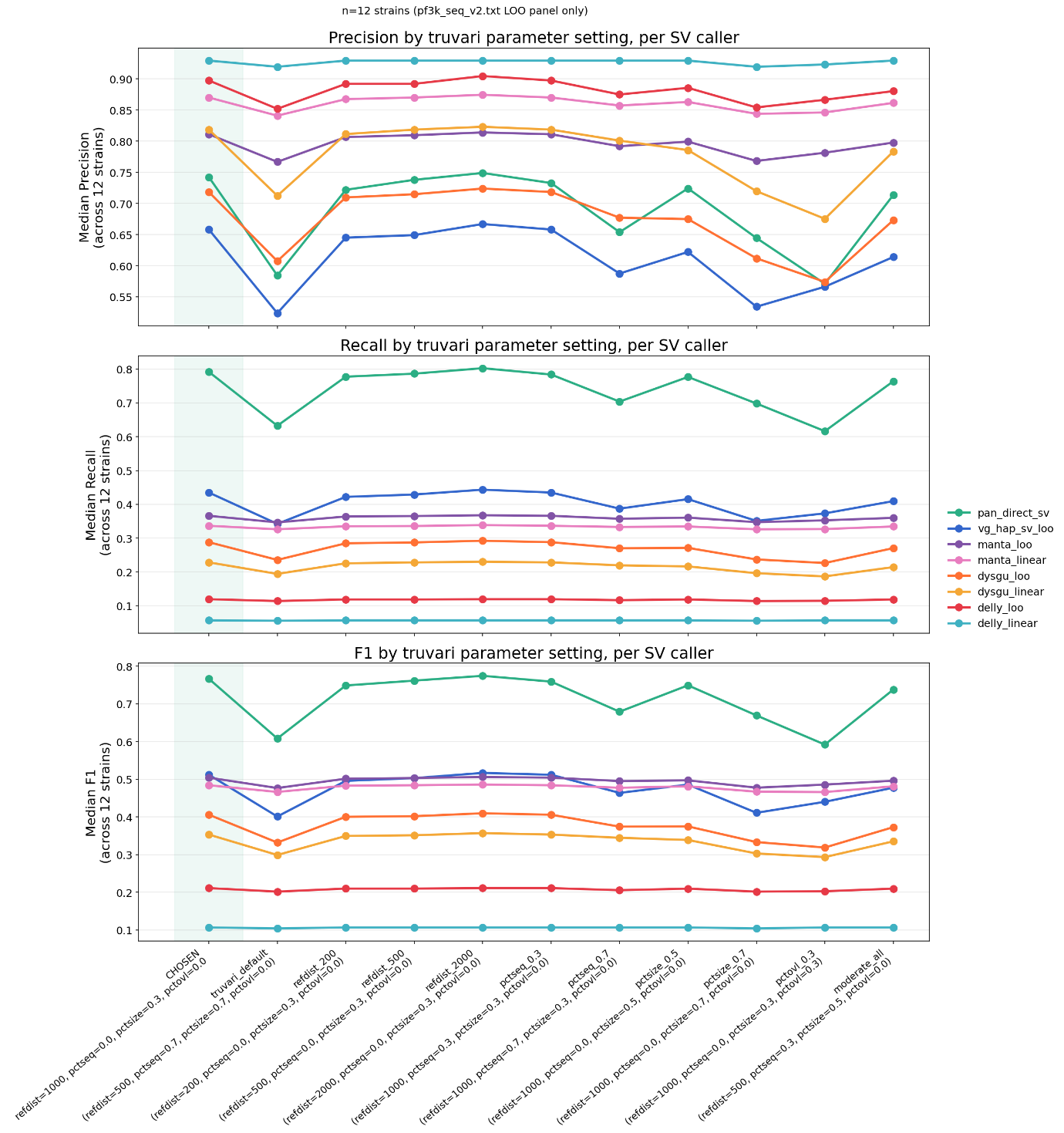

**Supplementary Figure 16. Sensitivity of SV benchmarking to Truvari matching parameters.** Median precision, recall and F-score of each SV caller across 12 leave-one-out strains, calculated within the core genome for SVs of 50 bp to 10 kb. Each x-axis entry is a Truvari parameter set. ‘Chosen’ is the setting used in the main analysis (--refdist 1000 --pctseq 0 --pctsize 0.3 --pctovl 0). "truvari_default" is refdist 500, pctseq 0.7, pctsize 0.7, pctovl 0. The refdist, pctseq, pctsize and pctovl entries change that single parameter from the chosen setting. "moderate_all" is refdist 500, pctseq 0.3, pctsize 0.5, pctovl 0. Lines connect settings for each caller to show the trend, not a continuous relationship. The same caller VCFs and truth set were used for every setting. Caller ranking is unchanged across all settings, whereas absolute values decrease as the criteria become stricter.
